# Radiotherapy Enhancement by Gold Nanocluster-functionalized Nanoliposomes Using Polychromatic Orthovoltage X-ray Irradiation

**DOI:** 10.64898/2025.12.04.692299

**Authors:** Nazareth Milagros Carigga Gutierrez, Eaazhisai Kandiah, Sofia Leo, Erwan Dupont, Fabrizio Johnson-Corrales, Marc-André Hograindleur, Sarvenaz Keshmiri, Hélène Elleaume, Benoit Busser, Romain Haudecoeur, Véronique Jacob, Anthony Vella, Véronique Josserand, Maxime Henry, Jean-Luc Coll, Amandine Hurbin, Xavier Le Guével, Mans Broekgaarden

**Affiliations:** Grenoble Alpes University, INSERM U 1209, CNRS UMR 5309, Institute for Advanced Biosciences, Site Santé, Allée des Alpes, 38700 La Tronche, France; European Synchrotron Radiation Facility, CM01 Beamline, 71 Avenue des Martyrs, 38000 Grenoble, France; Faculty of Pharmaceutical, Biochemical and Biotechnological Sciences, Catholic University of Santa María, Arequipa, Peru; Grenoble Alpes University, INSERM UA7, Synchrotron Radiation for Biomedicine, 38000 Grenoble, France; Grenoble Alpes University, CNRS, Department of Molecular Pharmacochemistry, 38000 Grenoble, France; Grenoble Alpes University, CNRS, IRD, Grenoble INP, Institute of Environmental Geosciences, 38000 Grenoble, France

**Author notes:** Corresponding author Mans Broekgaarden, Institute for Advanced Biosciences, Université de Grenoble-Alpes, Inserm U1209, CNRS UMR 5309, Allée des Alpes, 38700 La Tronche, France.

**Keywords:** Pancreatic cancer, liposomes, radiotherapy-responsive liposomes, gold nanoclusters, orthovoltage irradiation, reactive oxygen species, X-ray triggered drug release

## Abstract

Pancreatic ductal adenocarcinoma (PDAC) remains one of the most lethal cancers, as the most effective chemoradiation therapies achieve unsatisfactory outcomes while associated with high toxicity. Nanoliposomal drug delivery systems are widely used to improve chemotherapy safety, yet passive release of amphiphilic drugs may still associate with adverse toxicity. To address this, we previously developed nanoliposomes functionalized with hydrophobic gold nanoclusters, demonstrating radiocatalytic activity and enhanced chemoradiotherapy effects in 3D PDAC microtumors under synchrotron irradiation. In this study, gold nanocluster-functionalized nanoliposomes (AuLPs) were optimized and evaluated under 220 kVp orthovoltage X-ray irradiation, widely used in preclinical irradiation systems. AuLPs containing 0.2 mol% gold nanoclusters produced 1.5-fold more reactive oxygen species than unloaded liposomes. However, higher molar ratios were necessary to improve radiotherapy outcomes in 3D PDAC microtumor models following 4 and 8 Gy irradiation. Pharmacokinetics and biodistribution evaluations showed a modest increase in tumor gold content 24 hours post-injection in orthotopic PDAC models. Altogether, these results underscore the potential of gold-radiotherapy-responsive liposomes while highlighting critical formulation challenges, which must be resolved for full therapeutic potential.

## 1. Introduction

Pancreatic ductal adenocarcinoma (PDAC) remains one of the most lethal malignancies, with clinical failure frequently driven by early metastatic dissemination and pronounced resistance to therapeutic options. [1,2] Treatment resistance in PDAC is associated to multiple pathological barriers, which includes a dense desmoplastic stroma, elevated solid stress, high interstitial fluid pressures, and pronounced hypovascularity. These barriers significantly hinder the delivery and penetration of systemic therapies into the tumor core, [3,4] and may also play a role in radioresistance. [5] While the combination of chemotherapy and radiotherapy, often used as neoadjuvant or adjuvant therapy has improved outcomes for some patients with borderline resectable or locally advanced PDAC, [6–8] significant challenges remain. Notably, radiotherapy has been associated with a range of acute and late toxicities and potential damage to surrounding organs, as well as local recurrence, [9,10] thereby fueling efforts for more localized radiotherapy approaches.

Liposomes are valuable drug carriers that can reduce the toxicity of various chemotherapeutics. [11–13] Composed of a lipid bilayer that entraps an aqueous core, liposomes can encapsulate chemotherapeutics and increase their circulation times and tumor accumulation. [14] The latter occurs through the enhanced permeability and retention (EPR) effect, which refers to the preferential accumulation of nanoparticles in tumor tissue due to the leaky vasculature and poor lymphatic drainage commonly found in solid tumors. [15] Onivyde® (PEGylated liposomal irinotecan) illustrates this potential, achieving longer circulation, enhanced tumor localization, and greater efficacy of irinotecan at lower doses compared to the free drug. [16] Onivyde is now approved as a first-line treatment for metastatic PDAC in the NALIRIFOX regimen (with 5-fluorouracil, leucovorin, and oxaliplatin). [17] However, a major barrier to the efficacy of nanoparticle formulations in PDAC is their limited accumulation within tumors, largely due to the complex and fibrotic tumor microenvironment and limited blood perfusion due to high interstitial pressures. The dense extracellular matrix composed of a network of collagen and other stromal components impedes deep tissue penetration, often restricting nanoparticles to perivascular regions with limited quantities of drugs at the intended site of action. [18] Consequently, many nanotherapeutics fail due to dose-limiting toxicities in healthy tissues and insufficient delivery within the tumor mass.

A promising strategy to overcome these hurdles involves the use of externally triggered liposomes, which remain stable during circulation but release their payload upon activation by a stimulus. For example, light-responsive liposomes that enable photodynamic therapy have the ability to induce lipid membrane oxidation to facilitate controlled drug release, but also promote lysosomal permeabilization and prime the tumor microenvironment to enhance permeability. [19–25] However, their application is fundamentally limited by the shallow tissue penetration of visible and near-infrared light. [26] To address this, the use of ionizing radiation, specifically X-rays, has been proposed as an alternative trigger with the capacity to reach deep-seated tumors such as PDAC. The concept of radiotherapy-enhancing nanomedicines has garnered significant interest in recent years, particularly with the integration of high atomic number (high-Z) materials into nanoparticle platforms. These high-Z elements exhibit unique radio-physical properties that enable the amplification of local radiation dose through enhanced photoelectric absorption and the subsequent emission of secondary electrons, resulting in the generation of reactive oxygen species (ROS) in tissues, such as hydroxyl radicals (•OH) that can induce DNA damage and cell death. [27,28] In particular, gold nanoparticles have been extensively studied due to their high X-ray absorption coefficient and their ease of synthesis and surface properties modification, which enables precise control over the physico-chemical characteristics of the particles. [29,30] It has been described that the surface chemistry of gold nanoparticles can be optimized to improve their pharmacokinetics to increase tumor uptake and leverage its effects on radiotherapy enhancement. [30] Additionally, experiments conducted on mice carrying subcutaneous mammary carcinoma exhibited the radiosensitizing impact of 1.9 nm gold nanoparticles when exposed to 250 kVp X-rays, resulting in a significant survival enhancement compared to the treatment with X-rays alone. [31] Building on this approach, our group developed liposomes co-loaded with hydrophobic gold nanoclusters coated with dodecanethiol (AuDDT), and the photosensitizer benzoporphyrin derivative (BPD). Under synchrotron radiation, these Au+BPD co-loaded liposomes demonstrated significantly enhanced •OH production, particularly when irradiated with energies exceeding the K-edge of gold (80.7 keV). Additionally, calcein-loaded liposomes exhibited up to 30% release upon irradiation, indicating potential for controlled drug delivery. Notable therapeutic efficacy was observed in 3D pancreatic cancer models, where combining radiotherapy with oxaliplatin and Au+BPD liposomes resulted in reduced tumor viability and size. [32] Despite these promising findings, challenges remained. Notably, gold content within the liposomes was not quantified, and in vivo biodistribution studies failed to detect gold accumulation in tumors. Furthermore, the system relied on synchrotron radiation, which, while valuable for experimental studies, is not clinically applicable due to its limited availability and infrastructure requirements.

To bridge this gap, our current research focuses on evaluating the radiocatalytic potential of AuDDT-loaded liposomes under orthovoltage X-ray irradiation (220 kVp). A critical component of this work involved optimizing the encapsulation of AuDDT into liposomes. In addition to evaluating ROS production and drug release capacity, biodistribution studies were carried out in mice bearing orthotopic pancreatic tumors, using both fluorescence imaging and inductively coupled plasma mass spectrometry (ICP-MS) to confirm tumor targeting and pharmacokinetic profiles. The findings reveal the intriguing impact of X-ray source on radiotherapy enhancement by gold nanoclusters and highlight the limitations and challenges of this approach.

## 2. Results

### 3.1. Optimizing the preparation of AuDDT containing liposomes

To successfully encapsulate hydrophobic AuDDT nanoclusters within the lipid bilayer of liposomes, several parameters required systematic optimization. This led to the development of a specific preparation protocol for Au-loaded liposomes (AuLPs). The liposomal formulation consisted of DOPE:Cholesterol:DSPE-PEG in a 48:48:4 molar ratio.

One of the first parameters evaluated was the effect of sonication temperature following lipid-film hydration. Comparative analysis based on DLS measurements revealed that cold sonication (20 °C) yielded liposomes with significantly larger diameters (>200 nm), while hot sonication (60 °C) produced smaller vesicles (∼120 nm). This trend was maintained when increasing AuDDT loading from 0 to 2 mol% (Fig. 1A). Moreover, hot sonication led to a more homogeneous size distribution, as indicated by a lower PDI (Fig. 1B). Additionally, AuDDT encapsulation was semi-quantitatively evaluated using the intrinsic near-infrared fluorescence of the cluster. Cold sonication showed higher encapsulation efficacy compared to hot sonication (Fig. 1C). These findings illustrated the benefits of this approach for consistent liposome production. To improve liposomal uniformity, we performed extrusion through 0.2 µm filters. As expected, extrusion led to a substantial decrease in size and PDI, but at the expense of AuDDT encapsulation efficiency (Fig. 1D-F). This loss of AuDDT-loaded vesicles was attributed to their retention within the AlO matrix of the filters, as similar observations were made using cellulose filters in syringe-based extrusion systems.

**Figure 1.**
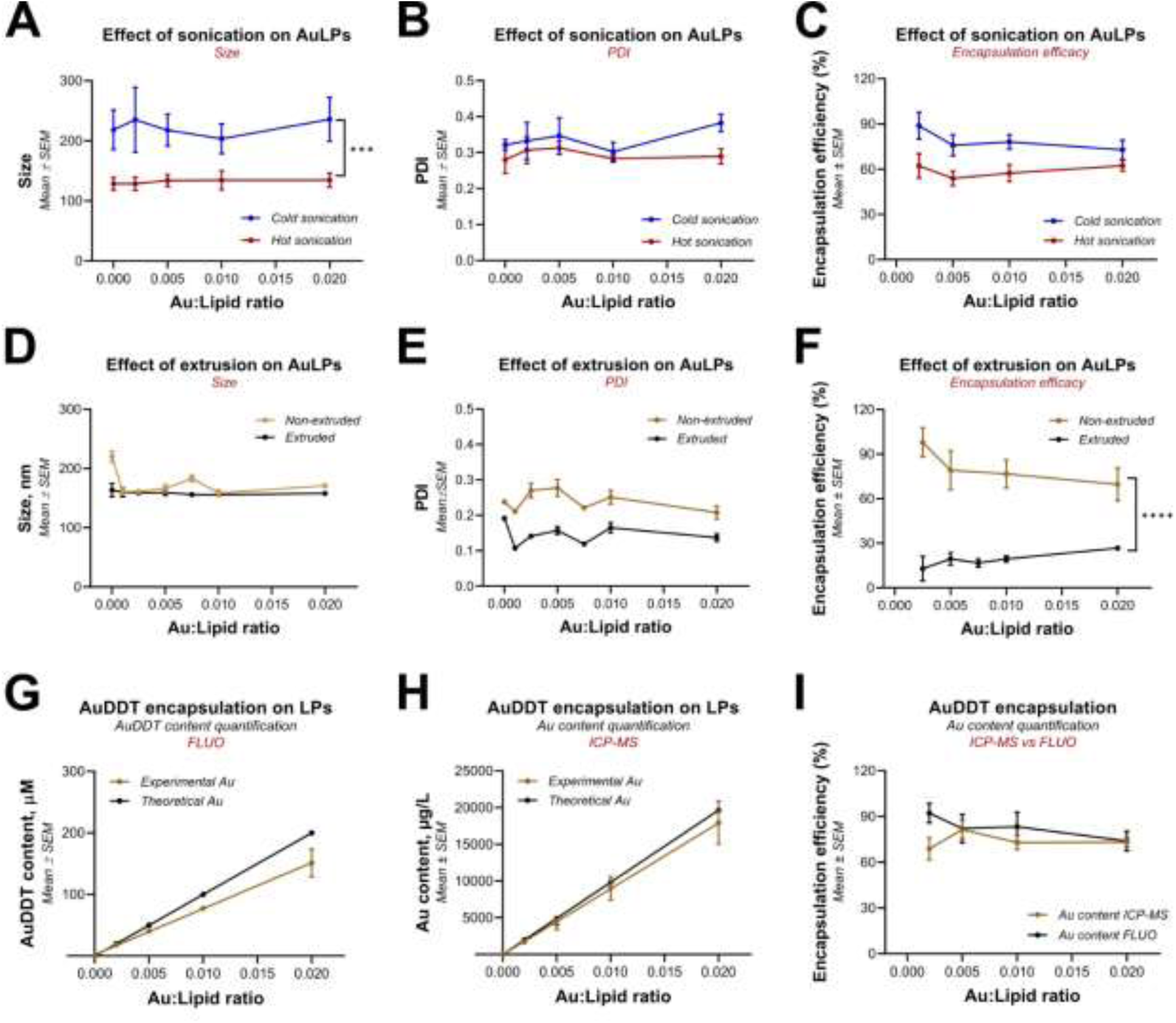
Engineering and optimizing AuDDT containing liposomes. (A) Effect of sonication temperature on liposome size, (B) polydispersity index (PDI), and (C) AuDDT encapsulation efficiency across formulations with increasing AuDDT content, cold sonication (20 °C) is shown in blue and hot sonication (60 °C) in red. (D) Effect of 0.2 µm extrusion on liposome size, (E) PDI and (F) AuDDT encapsulation efficiency across formulations with increasing AuDDT content (0 mol% to 2 mol%), non-extruded is shown in mustard and extruded in black. (G) Quantification of AuDDT content by SWIR-fluorescence and (H) ICP-MS, comparison between theoretical value shown in black and experimental value in mustard. (I) Comparison of encapsulation efficiencies determined by SWIR-fluorescence shown in black and ICP-MS in mustard. For A-C, the data represents n = 5-80 from seven independent batches. For D-F, the data represents n=5-25 from four independent batches. From G-I, the data represents n=5-86 from 20 independent batches. Linear regression fits were applied on G and H.

To evaluate the encapsulation efficacy of AuDDT within liposomes prepared by cold sonication, we compared near-infrared fluorescence emission and ICP-MS. Fluorescence intensities increased with the molar ratio of Au:Lipid, suggesting successful incorporation of gold clusters into the liposomes, producing values that closely matched the theoretical loading (Fig.1G). By ICP-MS, a clear linear correlation was observed between the initial estimated AuDDT input and the measured gold concentration in µg/L, confirming consistent encapsulation across formulations (Fig. 1H). The comparison of the two approaches revealed a high degree of concordance between the two methods, with encapsulation efficiencies converging at approximately 80% (Fig. 1I). These findings confirm that AuDDT was not only successfully encapsulated at the elemental level but also retained in its functional nanocluster form (Au25) within the liposomal membrane.

Having established a robust formulation process for AuLPs, we next assessed the influence of AuDDT concentration on their spectroscopic properties, particle size and colloidal stability. AuDDT nanoclusters, when incorporated into liposomes, displayed a bathochromic shift of the characteristic emission peak going from 1013 nm [33] to around 1050 nm (Fig. S1B and S1C). Such shifts were previous associated to the formation of assemblies or aggregates, [34] which may have occurred in the bilayer of the AuLPs. DLS analysis performed by backscatter mode revealed that all AuLPs, ranging from 0.2 to 2 mol%, exhibited a main size population centered around 100 nm (Fig. S1D). This demonstrates that AuLPs did not suffer from major vesicle enlargement after AuDDT loading. All formulations of AuLPs were colloidally stable, with average diameters ranging from 140 to 200 nm and low PDI values consistently below 0.3. No significant changes in either size or PDI were observed during a 4-week period (Fig. S1E-G).

### 3.2. Cryo-Microtomography confirms AuDDT integration within the liposomal bilayer of AuLPs

To confirm the spatial localization of AuDDT nanoclusters within liposomes, we performed cryo-EM on liposomal formulations containing 0, 0.5, and 2 mol% of AuDDT. Representative images reveal that, irrespective of AuDDT loading, all liposomes retained a spherical and primarily unilamellar morphology. Empty liposomes appeared uniformly translucent, with well-defined bilayers and no internal electron-dense features. In contrast, AuLPs exhibited discrete electron-dense circular spots embedded within their lipid bilayers, visible in the high-magnification images, as highlighted within red frames (Fig. 2A-B and S2A). The presence of electron-dense inclusions arises from the high atomic number of gold (Z = 79), which leads to strong elastic scattering of incident electrons, generating enhanced contrast in electron micrographs, [35] this is also consistent with our previous findings and other reports describing the incorporation of gold nanoclusters into lipid-based nanoparticles.[36–38,32] In our system, the presence of DOPE, a well-known fusogenic lipid, could have facilitated fusion events, which appeared to be further enhanced by the incorporation of AuDDT, as observed in some liposomal structures (Fig. 2 and Fig. S2). The consistent localization of AuDDT within the membrane, rather than in the aqueous core, further confirms its hydrophobic nature and preferential integration into the lipid bilayer.

**Figure 2.**
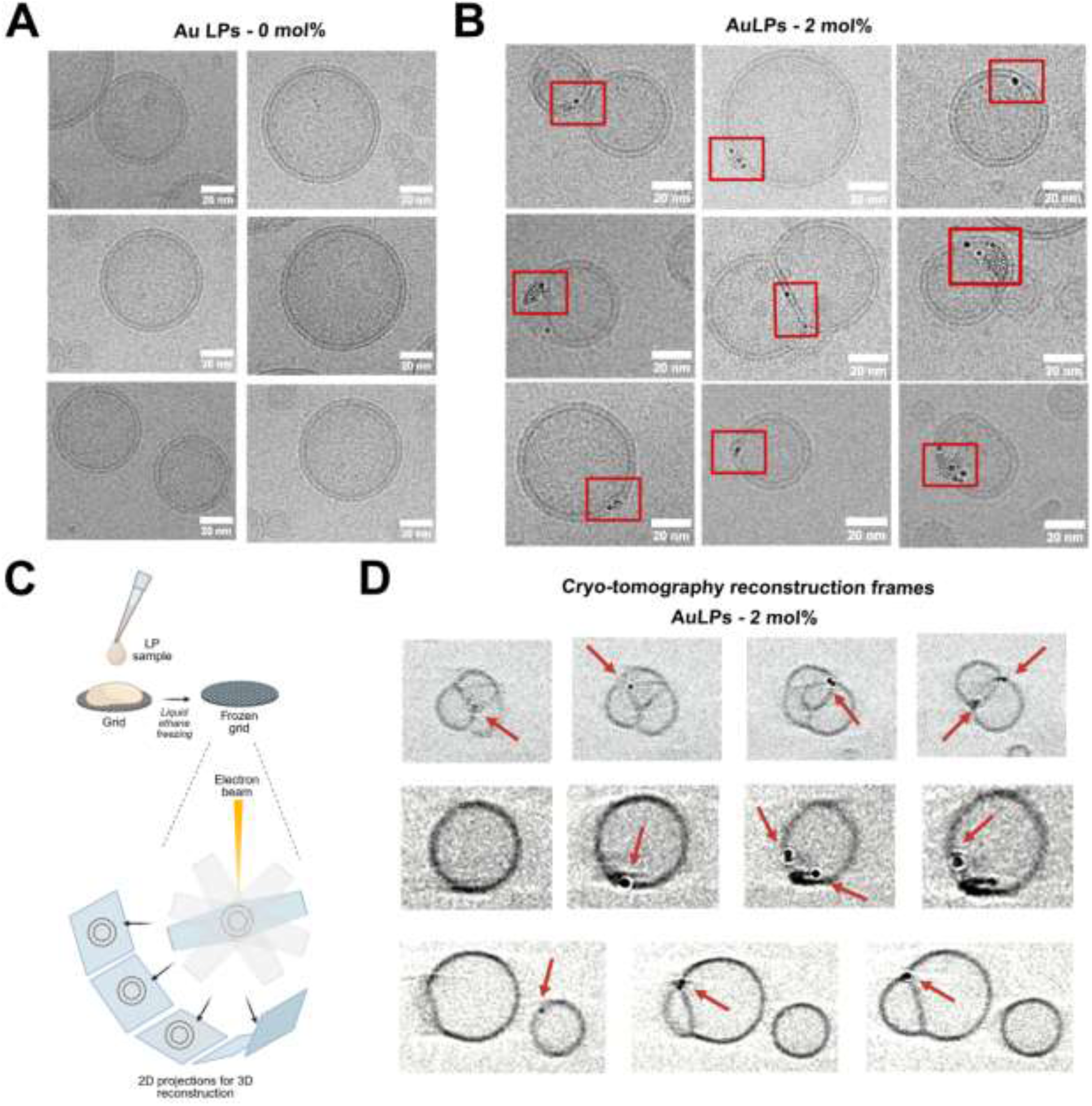
AuLPs morphological characterization through cryo-electron microscopy (cryo-EM). (A) Representative cryo-EM images of liposomes loaded with 0 or (B) 2 mol% of AuDDT (zoomed-in areas) distinct electron-dense spherical feature embedded in the lipid bilayer are identified as AuDDT nanoclusters, pointed out within a red frame (scale bar = 20nm). (C) Schematic representation of the cryo-electron tomography workflow, enabling spatial resolution AuDDT cluster localization. (D) Representative tomographic slices from 3D reconstructions of AuLPs containing 0.5 mol% AuDDT. Red arrows indicate AuDDT nanoclusters appearing and disappearing across different Z-planes.

Upon AuDDT incorporation, the membrane expanded to accommodate the nanoclusters, stretching the lipid bilayer as previously described. [32] These data suggest that the insertion of AuDDT imposes physical strain on the lipid bilayer, which undergoes lateral expansion to embed the clusters. This observation was further supported by a comprehensive morphological analysis of liposome structures across the full set of cryo-EM images. Despite the structural adaptations induced by AuDDT incorporation, the vesicle population preserved its overall morphological integrity. Lamellarity analysis revealed that approximately 80% of the vesicles were unilamellar, while only a minority (∼20%) exhibited multilamellar or fused configurations (Fig. S2B-D). This distribution was consistent across all tested formulations. Quantification and characterization of cluster occurrences revealed distinct populations differing in size and aggregation state, providing a semi-quantitative assessment of loading efficiency (Fig. S3). Cryo-electron tomography was performed to obtain three-dimensional reconstruction and more accurate assessment of the localization of the AuDDT. Analysis of sequential tomographic slices confirmed that several electron-dense features, were indeed embedded within the lipid bilayer but only became visible or fully resolved at specific Z-depths (Fig. 2D). This tomographic evidence strengthens our conclusion that AuDDT is consistently integrated into the liposomal membrane rather than adsorbed on the surface or dispersed in the surrounding medium.

### 3.3. Orthovoltage X-ray exposure (220 kVp) reveals ROS production but restricted drug release from AuLPs

We next evaluated how the AuLPs interacted with polychromatic orthovoltage X-ray irradiation. ROS generation occurred in a dose dependent manner. Non-loaded liposomes oxidized the APF probe at background levels, reflecting ROS generation from water radiolysis upon X-ray exposure. This was followed by liposomes containing the highest amount of AuDDT (2 mol%) (Fig. 3B). In contrast, liposomes with lower amounts of AuDDT (0.2 and 0.5mol%) generated significantly higher levels of ROS, suggesting that lower concentrations of gold nanoparticles may be adequate to enhance Au–X ray interactions. We selected the best performing AuLPs formulations, 0.2 and 0.5mol%, to investigate whether they could be used for radiotherapy-controlled chemotherapy delivery. To do so, the liposomes were loaded with a quenched fluorescent cargo (53 mM calcein), and its release was determined in response to X-ray stimulation. Minimal calcein release was observed across all X ray doses and AuDDT concentrations (Fig. 3C). Thus, despite the local ROS generation, the liposomal membranes remained largely intact, suggesting that the ROS were of insufficient in quantity or generated too far from the lipids, thereby failing to perturb bilayer integrity.

**Figure 3.**
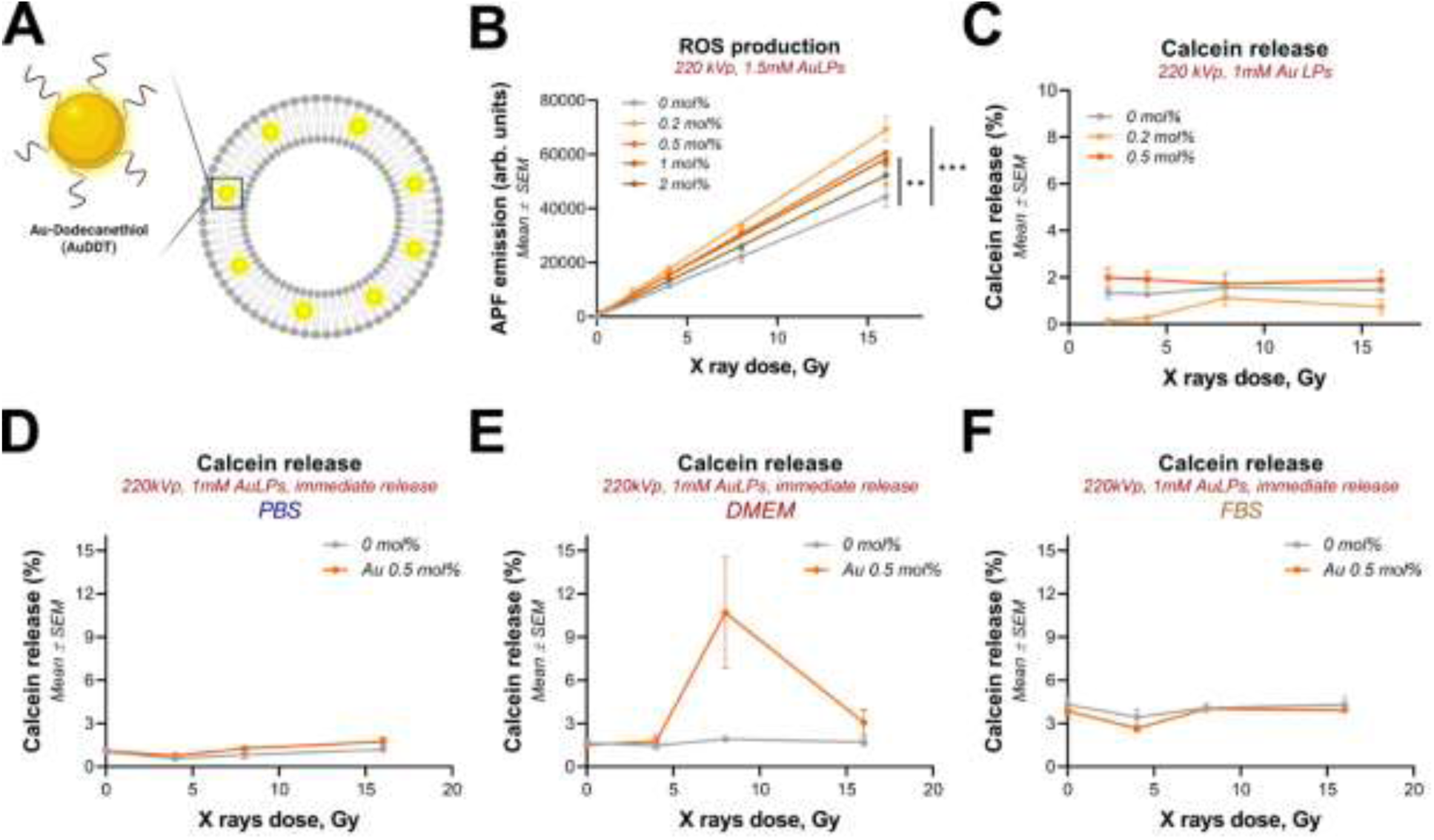
X-ray induced ROS production and triggered release from AuLPs. (A) Schematic representation of AuLPs with AuDDT integration in the lipid bilayer. (B) ROS production measured by APF oxidation under increasing doses of X-ray irradiation in different formulations of AuLPs with increasing AuDDT content (0-2mol%). (C) Immediate calcein release from AuLPs loaded with 0.2 mol or 0.5 mol% AuDDT after increasing doses of Xray irradiation. Immediate calcein release in different biological media: (D) PBS, (E) DMEM + 10% FBS and (F) FBS. For B-C, the data represent n = 18-25 from three independent technical repetitions, for D-F, the data represent n = 9-33 from three independent technical repetitions. Curve fitting was performed using a linear regression.

Previous studies have shown that gold nanoparticles and the photosensitizer BPD can act synergistically under high-energy X-ray irradiation (6 MeV). [39] We also observed this synergistic effect under synchrotron radiation, [32] where the synergy was hypothesized to come from the electron-donating capacity of porphyrins and their interaction with ionized gold atoms, possibly enhancing biochemical redox reactions. To investigate whether membrane sensitization could be enhanced by this synergy, we co-loaded BPD alongside AuDDT into the lipid bilayer (Fig. S4). However, neither radical production nor calcein release was observed to be elevated in the Au+BPD LPs (Fig. S4B-C), suggesting that the previously observed synergy at different X-ray energies (clinical megavoltage beams or synchrotron X-rays tuned above the gold K-edge) was not achieved at 220 kVp.

Finally, we assessed whether the release profile might depend on the surrounding medium. Three different solutions for ROS assay were compared: PBS (Phosphate Buffer Solution), FBS (Fetal Bovine Serum) and DMEM (Dulbecco’s Modified Eagle Medium) supplemented with 10% FBS. We observed an increase in calcein release of 10.1±17.18% in AuLPs at 0.5mol%, but only in DMEM at 8Gy (Fig. 3E). Regarding the release of calcein from AuLPs in PBS, we observed similar results as previous tests where the release was less than 3%. Finally, calcein release in FBS was very similar for loaded and non-loaded formulations showing a maximum release of 4.32±2.8%. This suggests that interactions between liposomes and media components may modulate membrane behavior under radiation, but further work is required to delineate these effects.

### 3.4. Increased AuDDT loading in AuLPs enhances cytotoxic effects and decreases PDAC 3D microtumor size

As the AuLPs produced a moderate yet measurable ROS production under X-ray irradiation, we next explored whether this could translate into therapeutic effects in a 3D in vitro pancreatic cancer model. In a first investigation, MIAPaCa-2, and PANC-1 microtumors grown on Matrigel were exposed to AuLPs (0.5 mol% AuDDT) at various liposomal concentrations ranging from 100 to 2000 µM, which were washed away following 8 h of exposure, followed by irradiation with increasing X-ray doses (4 and 16 Gy). While radiotherapy induced a notable decrease in microtumor viability in both microtumor models, the AuLPs did not exert any influence on the overall treatment efficiency (Fig. S5 and S6). These findings suggest that either the intracellular delivery of gold was insufficient to induce a biologically significant biological effect, or that, the degree of radiosensitization is limited by specific nanoscale energy deposition mechanism. [40] We therefore further explored whether increasing the AuDDT content in liposomes could enhance the therapeutic outcomes. To this end, we compared the standard formulation containing 0.5 mol% of AuDDT with formulations containing higher loadings of 2 mol%, 5 mol%, and 10 mol%. These AuLPs were tested at 2000 µM lipid concentration in 3D microtumors grown on Matrigel, where the washing step was omitted, and liposomes remained in contact with the cells during radiation. This setup partially mimics the EPR effect observed in vivo, where nanoparticles accumulate both within and around tumor tissues, allowing for both direct cellular uptake and indirect effects from the tumor microenvironment.

MIAPaCa-2 3D microtumors remained viable and morphologically healthy when incubated with AuLPs, even at the highest loading of AuDDT under no radiation conditions (Fig. 4A). Following irradiation at 4Gy, microtumor morphology showed evident structural alterations, accompanied by a homogeneous decrease in viability across all AuDDT loadings of AuLPs. At 8Gy, both viability and microtumor morphology were comparably affected across all formulations (Fig. 4B). Quantification of microtumor viability showed a slight, uniform decrease after 4 and 8 Gy irradiation, indicating that the effect was attributable solely to radiotherapy rather than AuLPs presence (Fig. 4C). With respect to size, non-irradiated microtumors remained stable, with the highest AuDDT loading (10 mol%) inducing only an 11.5% size reduction compared to non-loaded liposomes. Irradiation at 4Gy led to a reduction in mean tumor size across all formulations. This effect was more evident for formulations with >2 mol% loading, with tumor size decreasing by more than 28.6% compared to non-loaded liposomes. Notably, irradiation at 8Gy induced the greatest tumor size reduction, primarily due to radiation itself which decreased tumor size to 19.4±17% with non-loaded liposomes A marginally greater reduction was observed with 10 mol% AuLPs, lowering tumor size to 10.8±9% (Fig. 4D). Finally, analysis of residual viable disease (live fraction area) showed that radiotherapy, particularly at 4Gy, significantly reduced residual viable disease. Comparison with non-loaded liposomes indicated that even low loadings such as 0.5 mol% had an effect, with 5 mol% AuLPs appearing to be the most effective at 4Gy (Fig. 4E).

**Figure 4.**
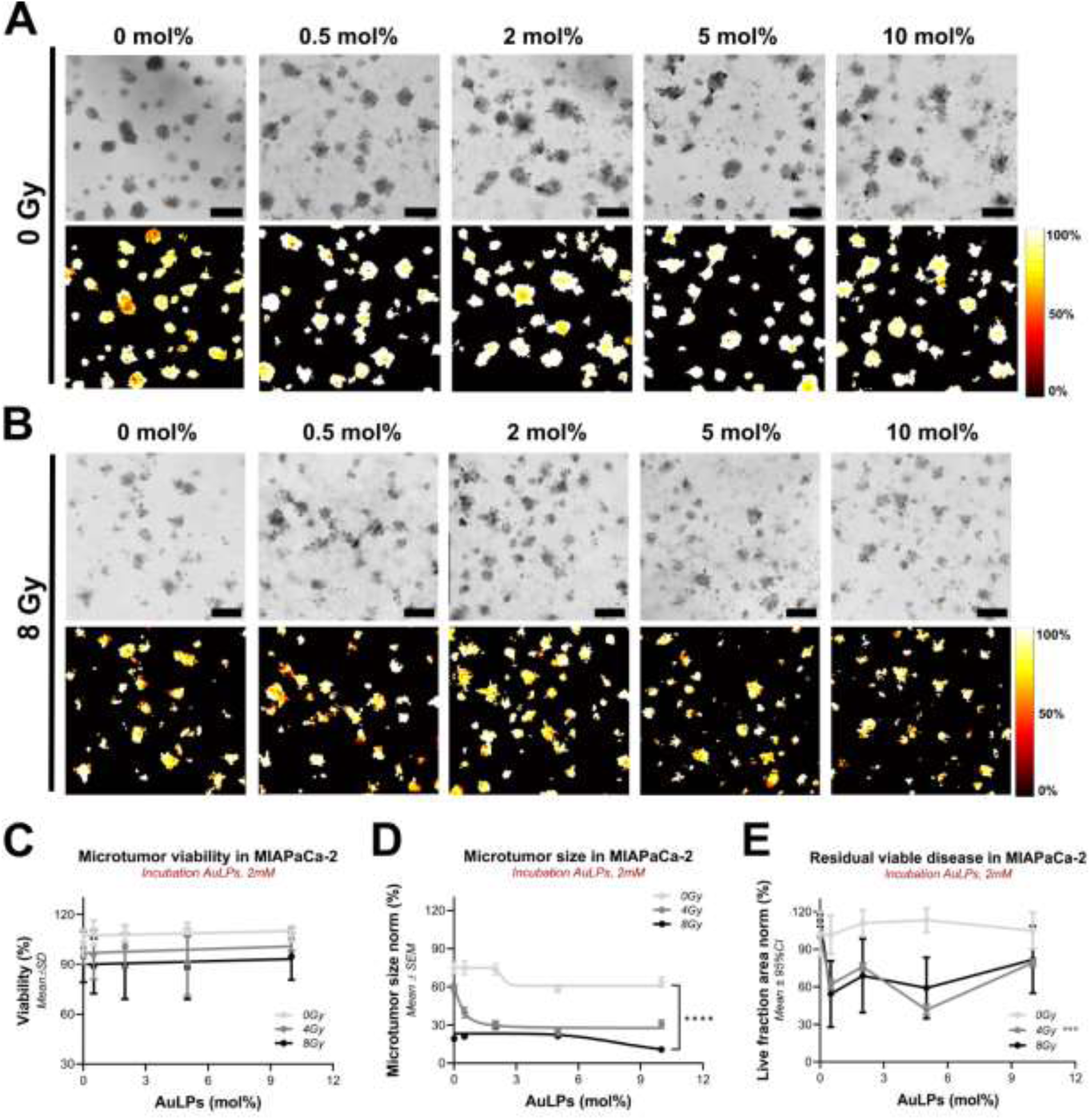
Effect of increased AuDDT content on AuLPs for evaluation of X-ray radiocatalytic effect on 3D MIAPaCa-2 microtumors. (A) Brightfield images and corresponding viability heatmaps of microtumors incubated with AuLPs with 0 to 10 mol% of AuDDT without irradiation (0 Gy), and at 8 Gy (B), scale bar 400 µm. (C) Quantification of microtumor viability in relation to AuDDT loaded on AuLPs for MIAPaCa-2 microtumors at different Xrays doses, 0, 4 and 8 Gy (Data was fitted with simple linear regression). (D) Quantification of normalized microtumor size in relation to AuDDT loaded on AuLPs for MIAPaCa-2 microtumors at different Xrays doses, 0, 4 and 8 Gy (curve was fitted for inhibitor vs normalized response-variable slope – four parameters). (E) Residual viable disease as the live fraction area remaining per condition, as AuDDT loaded on AuLPs (2mM) dose response for the different Xrays doses 0, 4 and 8 Gy. Data represent a sample size of n = 94-177.

Regarding PANC-1 microtumors, the AuLPs exhibited substantial toxicity and reduced microtumor size, effects that were further amplified by irradiation (Fig. 5A-B). Viability analysis revealed that even at 0 Gy, the highest of AuLPs loading (10 mol%) decreased viability by approximately 8% compared to non-loaded liposomes. This reduction was exacerbated under irradiation, with microtumor viability exhibiting a decreasing trend as a function of AuDDT content (Fig. 5C). In terms of size, a notable reduction was observed even in the absence of irradiation, where microtumors treated with 10 mol% AuLPs exhibiting the greatest shrinkage to 57.4% compared to the non-loaded liposomes. Under irradiation, this effect persisted and became more evident. In the absence of AuDDT, microtumor size was reduced to 38.4% and 28.2% at 4Gy and 8Gy, respectively. Increasing AuDDT content further enhanced this reduction, with 10 mol% AuLPs decreasing microtumor size to 15.1% at 4Gy and 14.7% at 8Gy (Fig. 5D). Quantification of the live fraction in PANC-1 indicated that AuLPs induced enhanced cytotoxicity in response to irradiation, even at low AuDDT concentrations such as 0.5 mol%. Formulations containing 2 mol% AuDDT exhibited the greatest reduction in viable disease (Fig. 5E).

**Figure 5.**
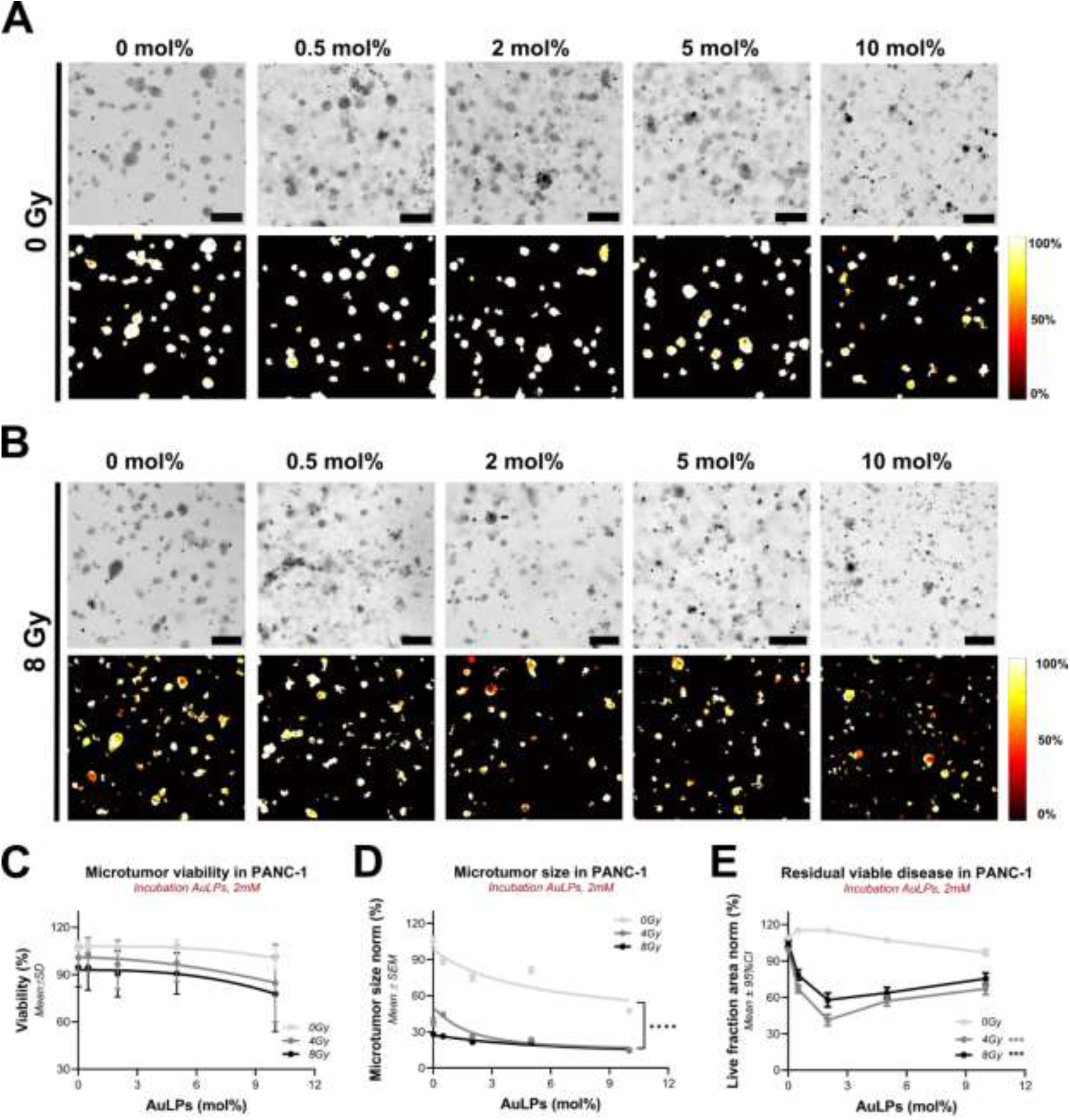
Effect of increased AuDDT content on AuLPs for evaluation of X-ray radiocatalytic effect on 3D PANC-1 microtumors. (A) Brightfield images and corresponding viability heatmaps of microtumors incubated with AuLPs with 0 to 10 mol% of AuDDT without irradiation (0 Gy), and at 8 Gy (B), scale bar 400µm. (C) Quantification of microtumor viability in relation to AuDDT loaded on AuLPs for PANC-1 microtumors at different Xrays doses, 0, 4 and 8 Gy (Data was fitted with simple linear regression). (D) Quantification of normalized microtumor size in relation to AuDDT loaded on AuLPs for PANC-1 microtumors at different Xrays doses, 0, 4 and 8 Gy (curve was fitted for inhibitor vs normalized response-variable slope – four parameters). (E) Residual viable disease as the live fraction area remaining per condition, as AuDDT loaded on AuLPs (2mM) dose response for the different Xrays doses 0, 4 and 8 Gy. Data represent a sample sizes of n = 109-348.

Overall, increasing AuDDT loading in AuLPs influenced therapeutic outcomes in a dose- and cell line–dependent manner. In MIAPaCa-2 microtumors, higher Au content did not substantially enhance cytotoxicity but promoted size reduction under irradiation, with 2–5 mol% formulations performing better than 10 mol%. In contrast, PANC-1 microtumors were more sensitive, showing baseline toxicity of the AuLPs, where the effects were further amplified by irradiation. Together, these findings indicate that while AuLPs can enhance radiotherapeutic responses, their efficacy is not linearly correlated with AuDDT loading, highlighting an optimal performance window between 2–5 mol%. Lastly, as liposomes containing more than 0.5mol% of AuDDT previously exhibited notable instability [32], we evaluated whether the liposomes containing 2, 5, and 10mol% AuDDT were stable when applying the optimized preparation protocol described in this study (Fig. 1). The results, displayed in Fig S7, surprisingly revealed that these liposomes remain stable over time, with minimal influence on liposome size, polydispersity, and particle concentration. However, after four weeks, the fluorescence emission of AuLPs markedly decrease compared to free AuDDT at equivalent concentrations, suggesting a loss of fluorescence activity and apparent reduction in estimated AuDDT encapsulation (∼60%)

### 3.5. Pharmacokinetics and biodistribution of AuLPs

Previous biodistribution studies for liposomes containing lipid-conjugated BPD (BPD-PC) and AuDDT at 0.5 mol% showed undetectable tumor accumulation of AuDDT and a divergent biodistribution profile compared to the BPD-PC-labelled liposomes. [32] However, it was unclear whether these findings were accurate given that the injected gold concentrations were low, and the detection sensitivity may have been insufficient. To resolve this, the biodistribution was re-evaluated by injecting a 4-fold higher liposome dose that contained 2 mol% of AuDDT rather than the 0.5 mol% AuDDT that was injected previously. Secondly, the injected liposome concentration was increased from 30 mg/mL in the previous experiment to 120 mg/mL in this experiment. A pharmacokinetic evaluation was done to investigate whether AuDDT was interacting with serum proteins. The pharmacokinetic profiles of both AuDDT and BPD-PC revealed similar blood circulation half-lives (Fig. 6A and 6B), with BPD-PC exhibiting a half-life of 1 h 50 min and AuDDT a slightly shorter half-life of 1 h 40 min. These comparable pharmacokinetics could suggest that BPD and AuDDT remain associated during systemic circulation in healthy mice.

**Figure 6.**
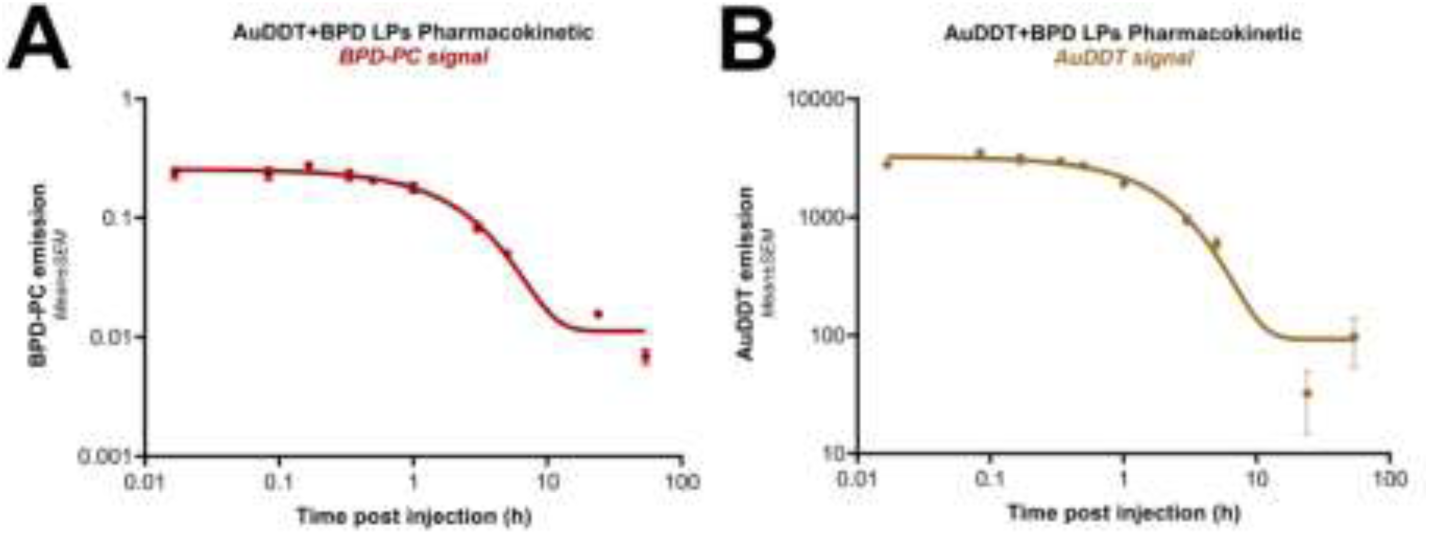
Pharmacokinetic distribution of AuDDT + BPD LPs on healthy mice. (A) Pharmacokinetic profile of BPD-PC signal collected over 54 hours in blood samples (in red) compared to control (in black). Pharmacokinetic profile AuDDT signal collected over 54 hours in blood samples (in mustard) compared to control (in black). The data represents n=3.

In contrast, the biodistribution profiles revealed a divergence in the organ level fate of these compounds. BPD-PC fluorescence was predominantly localized in the spleen, liver and kidneys, with accumulation at 3 h and 5 h post-injection and a marked decrease by 24 h, indicating progressive clearance, which can also be confirmed by full-body fluorescence imaging on mice (Fig. 7A and B). This aligns with previous studies where radiolabeled liposomes in the range of 160-200 nm have similar distribution profiles. [41,42] BPD-PC exhibited minimal accumulation and retention in the intestine, uterus, ovaries, adrenal glands, skin, muscle, and pancreas. Interestingly, the signal intensity in these organs was comparable to that observed in the tumor. Fluorescence from BPD-PC was still visibly detectable in *ex vivo* imaging up to 24 hours post-injection.

**Figure 7.**
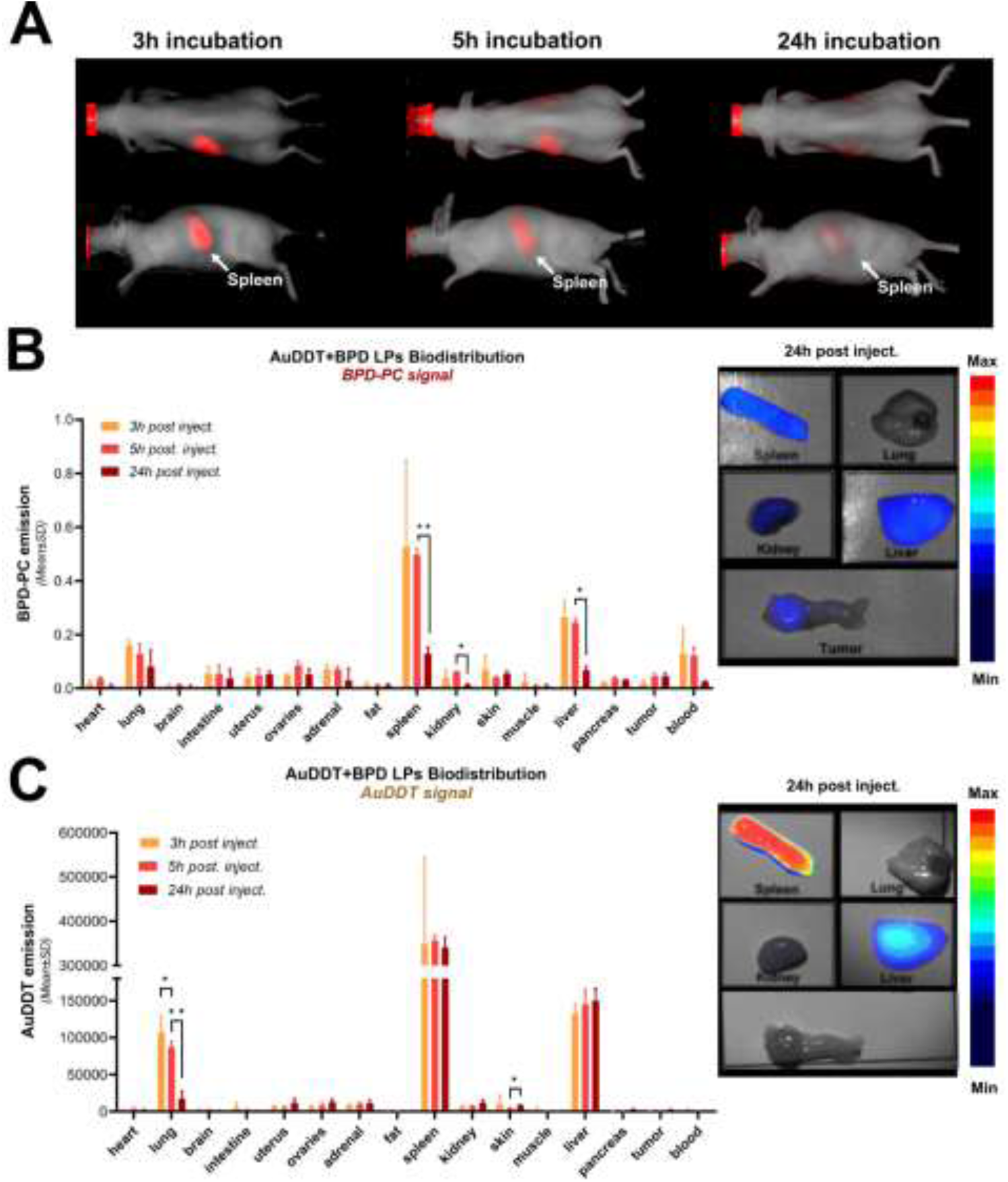
Biodistribution and pharmacokinetics Au+BPD LPs on mice. (A) Frontal and side views of a representative mouse injected with Au+BPD LPs at the different time points. (B) Biodistribution of BPD-PC fluorescence on mice bearing orthotopic pancreatic tumors, 3h, 5h and 24h post-injection, fluorescence signal on ex-vivo organs 24h post-injection are highlighted. (C) Biodistribution of AuDDT SWIR-fluorescence on mice bearing orthotopic pancreatic tumors, 3h, 5h and 24h post-injection, fluorescence signal on ex-vivo organs 24h post-injection is highlighted. The data represents n=9.

Compared to BPD-PC, AuDDT fluorescence also showed strong accumulation in the spleen and liver, but there was also a notable signal emitted from the lungs (Fig. 7C). In further contrast to BPD-PC, the organs displayed minimal clearance of AuDDT between 5 h and 24 h in the liver and spleen. These findings suggest that BPD-PC and AuDDT may follow distinct biodistribution or clearance pathways shortly after administration (Fig. 7B and 7C). These findings are in agreement with our previous study, in which AuDDT included at a 0.5mol% liposomal composition was cleared from spleen and kidneys 24 hours post injection. [32] The data suggests that AuDDT escapes from the liposome upon absorption while circulating in the organs, with limiting biliary excretion and, consequentially, long retention in the spleen and liver. [43,44]

The discrepancy with the pharmacokinetic data may hint towards the inaccuracy of determining AuDDT concentrations by SWIR fluorescence imaging when performed in presence of BPD-PC. Therefore, ICP-MS was employed for tissue analysis at 3, 5, and 24 h post-administration. The ICP-MS data corroborated the SWIR imaging findings, demonstrating persistent gold retention in liver and spleen through 24h, with emerging tumor accumulation over time (Fig. 8A-C). To further confirm gold accumulation in tissues, laser-induced breakdown spectroscopy (LIBS) was conducted on adjacent tissue sections corresponding to the highest gold levels detected by ICP-MS. Quantification of gold and phosphate (general tissue marker) was performed in spleen, liver, and tumor tissues (Fig. S8). Consistent with ICP-MS results, LIBS confirmed the presence of gold in tumor sections at 24 h post-injection, though at low intensities. To enable more precise comparison between samples, the S-value, a normalized measured of Au signal per unit tissue area, was employed. This quantitative metric has previously demonstrated reliable correlation between LIBS and ICP-MS measurements for titanium (Ti) quantification. [45] These values were observed to increase over time, measuring 1.88×10^4^, 3.03×10^4^ and 1.74×10^5^ for tumor tissues at 3, 5 and 24 hours, respectively (Fig. 8D). Adjacent tissue sections stained with hematoxylin and eosin (H&E) were used to assess tissue morphology and confirm tissue architecture of areas analyzed by LIBS. This quantification further confirms the SWIR fluorescence and ICP-MS measurements, where gold content increases over time.

**Figure 8.**
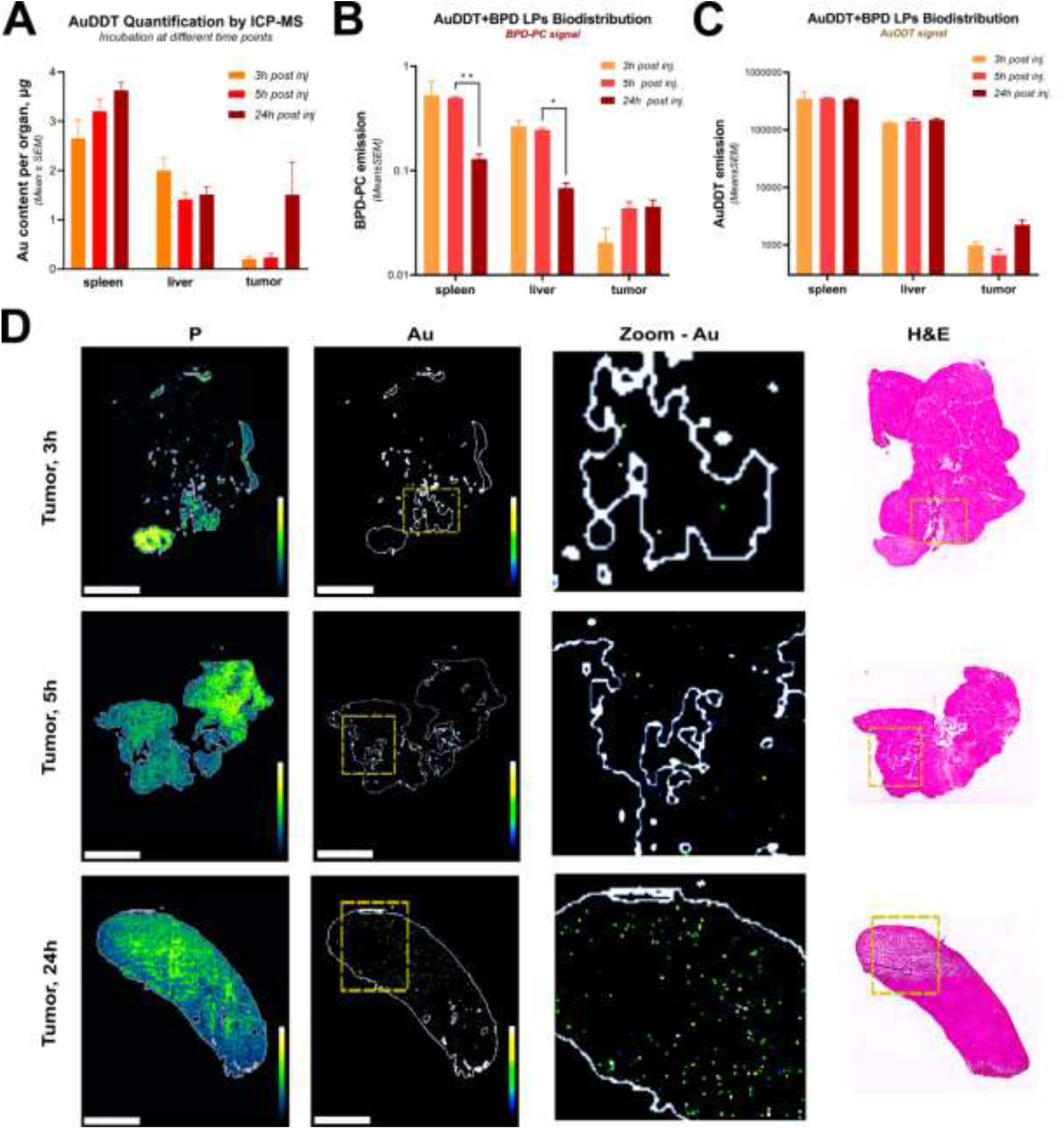
Quantitative analysis of AuLPs distribution and histological evaluation in tumor-bearing mice. (A) ICP-MS quantification of Au content in spleen, liver and tumor, at 3h, 5h and 24h post liposome injection. (B) BPD fluorescence signal in spleen, liver and tumor, at 3h, 5h and 24h post liposome injection. AuDDT signal intensity in spleen, liver and tumor, at 3h, 5h and 24h post liposome injection. Data represents n=9 from three technical repetitions. (D) Elemental mapping of orthotopic PANC-1 tumor sections by LIBS for quantification of phosphorus (P) and gold (Au) on Au+BPD treated mice, depicted are tumor tissue, at 3h, 5h and 24h post liposome injection and representative H&E staining of the tissue slide. Scale bar, 2 mm.

However, the amount of gold detected in tumor tissues by ICP-MS and confirmed by LIBS at 24 hours post-injection is relatively low. While gold accumulation in tumors increases over time, the concentrations measured may still be below or at the lower threshold required to achieve effective radiosensitization effects. Furthermore, the discrepancy between AuDDT and BPD-PC fluorescence signal highlights the challenges in quantifying and following AuLPs concentrations in vivo. Additional strategies, such as optimizing the formulation and enhancing AuDDT retention, may be required to achieve therapeutically relevant gold concentrations in tumors.

## 3. Discussion

This study investigated the optimization of liposome formulation containing of hydrophobic gold nanoclusters (AuDDT), which previously demonstrated to be radiotherapy-responsive nanocarriers under monochromatic synchrotron radiation (81.7 keV). This study builds upon those findings and presents an optimized preparation procedure that yields stable nanoliposomes capable of containing significantly elevated amounts of up to 10mol% AuDDT, compared to our previous study. While monochromatic synchrotron radiation is valuable for fundamental studies, it is not widely accessible and has limited relevance for preclinical studies that frequently utilize polychromatic orthovoltage X-rays. This study therefore explores the interaction of our optimized AuLPs formulations using 220 kVp X-ray irradiation. While the AuLPs exhibited ROS generation under orthovoltage irradiation, the fold-change compared to empty liposomes was 1.68, which was significantly lower compared to ∼8 upon 81.7 keV irradiation. [32] This reduced radiocatalytic efficiency also translated to reduced efficiency of cargo release from the AuLPs. Whereas the release of 21% of the entrapped calcein was reported at 81.7 keV, only 1.7% was released under similar conditions using 220kVp. Interestingly, AuLPs improved radiotherapy outcomes in 3D culture models of pancreatic cancer, which exceeded expectations considering their modest radiocatalytic properties under 220kVp. These finding may suggest that the AuLPs could induce radiosensitization via biological mechanisms rather than physical, which could merit further investigation. Lastly, while previous biodistribution and tumor accumulation studies where somewhat inconclusive, the more compelling approach applied here confirms our previous indications: the AuLPs do not stay intact upon systemic injection and produce different biodistribution profiles for lipid-conjugated dyes and AuDDT. The modest tumor accumulation of AuDDT that results remain discouraging for further radiotherapeutic studies in vivo using orthotopic pancreatic cancer models.

Considering the optimization of the preparation procedure of the AuLPs, we made several interesting observations. Chloroform was employed as the phospholipid preparation solvent, as it has been reported to facilitate compound incorporation into lipid membranes. In line with these reports, its use in our system enabled efficient AuDDT encapsulation, likely by reducing bilayer viscosity and promoting nanoparticle integration at the membrane - water interface. Notably, chloroform was fully evaporated during lipid film formation and not retained in the final liposome solution [46,47] Regarding sonication temperature, we observed that sonication at approximately 60°C produced a stable, homogeneous liposome population with relatively small sizes (<150 nm), whereas sonication at lower temperatures around 30°C favored a better encapsulation of AuDDT. This observation aligns with previous studies on lecithin liposomes demonstrating that increasing the sonication temperature from 50°C to 70°C reduces both liposome size and polydispersity index, likely due to increased fluidity of the lipid bilayer enhancing the efficiency of ultrasonic disruption. [48] Conversely, sonication at ∼30°C has been reported to yield more polymorphic structures with unincorporated lipids. [49] At this temperature it appears that the reduced bilayer fluidity preserves better hydrophobic cargo limiting leakage during sonication. Extrusion of AuLPs led to a substantial loss of AuDDT, consistent with previous observations that size-reduction via extrusion can compromise drug encapsulation. This loss likely arises from a combination of factors related to the mechanical stress imposed during extrusion. First, the high shear forces and pressured experienced by liposomes as they are forced through nanoscale filters can disrupt membrane integrity, causing leakage of hydrophobic cargo such as AuDDT. Additionally, some liposomes or aggregated complexes may become physically trapped or adhere within the filter membranes, particularly when filter pore size approaches the liposome size, further reducing observed encapsulation. The extrusion process thus highlights an inherent trade-off between achieving size homogeneity and preserving encapsulation efficiency. [50] Moreover, comparative analysis of different extrusion devices further supports that differences in flow rates and shear stress correlate with greater leakage and membrane disruption. [51] The understanding of these mechanisms is crucial for optimizing liposome uniformity and cargo retention during preparation.

Cryo-EM confirmed our previous findings of AuDDT inclusion into the lipid bilayers. AuDDT inclusion in the lipid bilayer appeared to cause lateral expansions between the membrane leaflets to accommodate the clusters. Increasing AuDDT content from 0.5 [32] mol% to 2 mol% appeared to exacerbate these effects, as higher loadings likely induced greater bilayer strain and partial leakage of AuDDT, consistent with the instability observed in biodistribution studies. Such behavior aligns with previous reports describing the interaction between gold nanoclusters and lipid bilayers, where the inclusion of hydrophobic gold nanoclusters did not compromise membrane stability but occasionally promoted vesicle fusion or increased lamellarity, depending on the formulation and preparation conditions. For example, in pure DOPC vesicle membranes, 4.1 nm Au nanoclusters impaired vesicle formation and modified vesicle structures by creating lipid assemblies, [52] it has also been reported that interactions with the lipid bilayer are dependent on the size and concentration of the nanoparticles. [53,54]

Concerning the radiocatalytic properties of the AuLPs during 220kVp irradiation, it appears that the spatial distribution of gold nanoparticles plays a critical role in mediating radiation enhancement. *McMahon et al*. revealed that when X-rays interact with gold nanoparticle systems, only a small fraction of the gold atoms within the nanoparticles are ionized. This results in a heterogeneous or clustered dose distribution at the nanoscale, because not all gold nanoparticles absorb X-ray energy equally or simultaneously. Following ionization, gold atoms release both high-energy photoelectrons, which travel relatively long distances, and low-energy Auger electrons, which have a very short range. While photoelectrons contribute to overall energy deposition, it is the Auger electrons that primarily drive localized dose enhancement. [40] The location of the ionization event within the gold nanoparticle also plays a pivotal role on its radiocatalytic effectiveness, especially when AuDDT aggregates were observed within the liposome bilayers. Due to the relatively short range of low energy Auger electrons, ionization events occurring deep inside larger nanoparticles are unlikely to contribute to dose enhancement, as these electrons are re-absorbed by surrounding gold atoms within the nanoparticle where they are generated. In such cases, these low-energy electrons do not escape to the surrounding medium. In contrast, ionization events occurring near the nanoparticle surface are more effective at emitting low-energy electrons into the surrounding medium, thereby promoting the generation of ROS. When nanoparticles are aggregated at high concentrations, most gold atoms are in the core of the cluster, limiting surface events. In contrast, at low concentrations, gold nanoparticles are better dispersed within the medium, allowing for more surface interactions and resulting in more efficient ROS generation. We therefore suspect that the AuLPs with high molar ratios of AuDDT exhibit quenched radiocatalytic activity due to aggregation.

Although AuLPs exhibited measurable ROS generation under 220 kVp X-rays, this was insufficient to induce significant X-ray–triggered release of liposomal cargoes in PBS. Intriguingly, partial release was detected in DMEM, suggesting that serum proteins or amino acid components could promote lipid oxidation during radiotherapy. Such behavior has been reported for other liposomal delivery systems, where release of cargo was 5-times faster in DMEM than in PBS, which was was attributed to the presence several factors such as albumin, forming a corona that modifies colloidal stability and liposomal permeability. [55,56] These results indicate that although radiocatalytic activity occurs under orthovoltage irradiation, achieving effective membrane oxidation likely requires lipid compositions with higher susceptibility to oxidation or incorporation of more reactive radiocatalytic agents to sufficiently compromise membrane integrity. A peculiar finding was that the release reproducibly occurred at a dose of 8Gy, whereas lower and higher doses did not achieve notable cargo release. One can speculate that more complex radiochemical reactions are the underlying cause of this observation, such as a threshold dose required to augment ROS generation, and the consumption of oxygen or redox intermediates within the culture medium being a limiting factor at higher doses. Interestingly, a radiotherapy enhancement effect on pancreatic microtumors in vitro was observed only when the washing step was omitted and AuDDT loading exceeded 2 mol%. This effect was more pronounced in PANC-1 than in MIAPaCa-2 cells. The prolonged incubation of liposomes with cells before and during irradiation likely contributed to a combined radiosensitizing response, potentially involving both physical and biological radiosensitizing mechanisms. Indeed, gold nanoparticles have been reported to enhance ROS generation through NADPH oxidase activation, mitochondrial dysfunction, and oxidation of intracellular antioxidants, suggesting that both physical and biological factors may underlie the observed effects, warranting further investigation. [57,58]

Motivated by our previous results and others on the synergy between gold nanoparticles and the photosensitizer BPD under distinct irradiation sources (MeV and synchrotron radiation), [39,32] AuDDT+BPD liposomes were also tested. However, at 220 kVp no synergistic interaction was observed, neither for ROS production no for calcein release. It is possible, that at this energy, gold ionization is insufficient to promote its interaction with BPD, or that the resulting ROS levels remain below the threshold required to induce membrane rupture. Alternatively, the dose rate differences of 3.06 Gy min^-1^ (220 kVp), 52.8 Gy min^-1^ (monochromatic synchrotron radiation), and 1 to 10 Gy min^-1^ (6 MVp linear accelerator) could also account for the discrepancy in our findings. Moreover, their radiocatalytic effect in 3D spheroids did not exhibit any significant enhancement when using Au+BPD co-loaded liposomes.

Concerning the biodistribution and tumor accumulation, our previous results indicated that AuDDT loadings of 0.5 mol% were insufficient to produce a detectable gold signal in tumors. This poor accumulation, along with a divergent biodistribution pattern compared to the liposomal (BPD-PC) signal, posed a major limitation for further in vivo evaluation. In the present study, AuDDT loading was increased to 2 mol%, aiming to enhance tumor accumulation and improve detection sensitivity. Although a slight increase in gold signal was detected 24 hours post-injection, the overall intratumoral concentration remained low and inadequate to achieve effective radiosensitization. Previous studies have demonstrated that radiation dose enhancement requires high-Z element concentrations in the range of ∼10-500µg per gram of tumor tissue, substantially above the gold concentrations observed in out cancer tissues (<5µg/gr of tissue).[59] Moreover, the observed differences between the biodistribution profiles from AuDDT and BPD-PC suggest that AuDDT also appeared to escape into serum proteins, as previously hypothesized, [32] and the increased AuDDT content may have further compromised membrane stability. This aspect is critical to consider for formulation optimization, as membrane stability strongly influences the retention of encapsulated compounds, tissue penetration, and interactions with serum proteins, parameters that can be modulated by adjusting cholesterol content or incorporating lipids with higher phase transition temperatures. [60,61]

The results of this study reveal fundamental insights into the engineering of X-ray-responsive liposomes suitable for orthovoltage radiation sources. Our findings indicate that radiotherapeutic efficacy can be significantly enhanced by incorporating radiocatalytic elements, particularly when these elements are embedded within liposomal membranes. Nonetheless, several limitations inherent to the current generation of AuLPs must be addressed to reach their full potential. Key challenges include increasing their radiocatalytic capacity to induce efficient X-ray-triggered release, which could be achieved by loading higher concentrations of AuDDT or systematically optimizing the liposomal membrane composition. For instance, the successful loading of gold clusters into liposome cores as well as lipid-conjugated gold clusters has been demonstrated, offering promising avenues for increased payload stability and responsiveness. [62,63] Moreover, AuDDT nanoclusters are coated with dodecanethiol ligands, a class of thiols well-known for their potent antioxidant properties. Thiols can act as ROS scavengers due to their sulfhydryl (-SH) groups, which donate hydrogen atoms to neutralize free radicals. This inherent antioxidant capacity of the thiol coating may represent a major limitation for AuLPs, as it can partially quench ROS generation and thus reduce the efficiency of radiation-induced cytotoxic effects. [64,65] In addition, incorporating higher amounts of (poly)unsaturated phospholipids could improve responsiveness to oxidative stress, as previously demonstrated in light-triggered release systems. [24,66] Finally, the inclusion and modulation of other components such as cholesterol and DSPE-PEG are also likely to have a significant impact on release behavior and stability, warranting further investigation. [67,68] Another critical aspect is improving tumor accumulation while minimizing AuDDT leakage in circulation. This may be addressed through enhanced gold loading and potential crosslinking strategies of gold nanoclusters to liposomal membranes, thereby improving retention and systemic stability. [69] However, when co-loading chemotherapeutics, it is essential to balance membrane rigidity to maintain gold retention without compromising drug encapsulation efficiency. Optimizing these parameters is crucial to developing liposomal platforms capable of reliable, X-ray-triggered delivery for clinical translation.

## 4. Conclusion

This work establishes the development and evaluation of orthovoltage-responsive AuDDT liposomes designed for radiocatalytic enhancement in pancreatic cancer. Although their radiotherapy enhancement effect was modest, the study identifies key formulation parameters governing the interplay between gold distribution, lipid composition, and biological stability. These findings underscore the necessity to refine liposomal design to improve gold retention and radiocatalytic accessibility within the membrane. Enhancing tumor accumulation and minimizing gold leaching remain critical challenges that can be addressed through optimized AuDDT loading and potential crosslinking of gold nanoclusters to liposomal membranes. Successfully balancing membrane rigidity to retain gold without compromising chemotherapeutic encapsulation will be essential for developing an effective X-ray-triggered drug delivery platform with strong translational potential in pancreatic ductal adenocarcinoma treatment. This integrated approach, aligned with ongoing innovations in radiocontrol strategies, paves the way toward realizing clinically impactful nanotherapeutics for radiotherapy augmentation.

## 5. Materials and Methods

### Chemicals and reagents

The lipids 1,2-distearoyl-sn-glycero-3-phosphocholine (DSPC), 1,2-dioleoyl-sn-glycero-3-phosphoethanolamine (DOPE), 1,2-distearoyl-sn-glycero-3-phosphoethanolamine-N- [methoxy(polyethylene glycol)-2000 (DSPE-PEG), were purchased from Avanti Polar Lipids (Alabaster AL, USA), cholesterol was from Sigma-Aldrich (Merck, Darmstad, Germany), and 1-stearoyl-2-hydroxy-sn-glycero-3-phosphocholine (18:0 Lyso PC) was purchased from Lipoid (Ludwigshafen, Germany). Lipid stock solutions were prepared in chloroform and stored under nitrogen gas at −20°C. Hydrophobic gold nanoclusters stabilized using dodecanethiol (AuDDT) were prepared as described previously [70,71]. Briefly, 46.4 mg (93.75 µmol) AuClPPh_3_ were dispersed in ethanol followed by addition of 81.5 mg (0.94 µmol) tert-butylamine-borane and fast stirring for 45 min. The solution changed from orange red to black color after 5 min. Then 12 µL of DDT was added and the mixture was left to stir for another 3 h. The dark solution was then filtered 3 times with filters at 3 kDa molecular weight cut-off to remove unreactive species and kept refrigerated. AuDDT was suspended in chloroform at a concentration of 3.5 mg Au per mL. Benzoporphyrin derivative (BPD) was purchased from Sigma-Aldrich (Darmstadt, Germany). Calcein, benzoporphyrin derivative (>94%), Sephadex G50 fine (Cytiva, Marlborough, MA, USA), and nitrogen gas were acquired from Sigma-Aldrich (Merck). N-(3-dimethylaminopropyl)- N′-ethylcarbodiimide (EDC), 4-dimethylaminopyridine (DMAP,) and N,N-diisopropylethylamine (DIPEA 0.742 g/mL) were purchased from Sigma Aldrich. Solvents used for reactions and purifications including dichloromethane (DCM) and methanol (MeOH) were purchased from VWR (Rosny-sous-Bois, France).

### Synthesis and characterization of lipid-conjugated benzoporphyrin-derivative (BPD-PC)

18:0 BPD-phosphocholine (BPD-PC) was synthesized via an adaptation of previous procedures [20]. In a two necked flask under an argon atmosphere, 1-stearoyl-2-hydroxy-sn-glycero-3-phosphocholine (24.77 mg, 0.047 mmol) was dissolved in 5 mL of DCM with the aid of sonication. Subsequently BPD was introduced to the stirring mixture (17 mg, 0.024 mmol) together with EDC · HCl (3.67 mg, 0.024 mmol) and the DMAP (14.45 mg, 0.118 mmol). Finally, DIPEA was added into the solution in a drop-wise fashion (41.2 μL, 0.237 mmol). The reaction was stirred for 72 hours and monitored on daily basis by TLC (Silica gel 60 F254, aluminum sheets, Supelco, Sigma-Aldrich) and UV-Vis spectroscopy (UV-1800 Shimadzu, Kyoto, Japan). The reaction products were purified by chromatography using diol-silica (110 Å pore size, 40-75 μm, Sigma-Aldrich, Germany) as stationary phase and modulating DCM and MeOH as mobile phase. Pure products were characterized through ^1^H-NMR (Bruker Ultrashield 400TM, Billerica, USA) and MALDI-TOF (Bruker microflex®, Billerica, USA) (Fig. S9).

### Liposome preparation and characterization

Liposomes were prepared using the lipid-film hydration method, where stock solutions were prepared in chloroform for cholesterol (50 mM), DOPE (50mM) and DSPE-PEG 2000 (10m M) were mixed at molar ratios of 48:48:4. BPD (2.77 mM in CHCl_3_) was added to the lipid mixture at a lipid:BPD ratio of 0.008, AuDDT (17.8 mM) was included at increasing lipid:AuDDT ratio from 0.002 to 0.02 (0.2 to 2 mol%) for loading and characterization purposes. Lipid films supplemented with AuDDT and/or BPD were prepared using vacuum centrifugation (Genevac miVac centrifugal concentrator, Ipswich, UK, 40°C, 4 h minimum), hydrated in phosphate-buffered saline (PBS), and sonicated at 37 kHz (Elmasonic p30, Elma Schmidbauer GmbH, Singen, Germany), with the sonication temperature being either 20°C (cold sonication) or 60 °C (hot sonication) until a clear solution was obtained. Similarly, parameters such as stock organic solvent, extrusion, and sonication time were evaluated. When liposomes were extruded, inorganic membrane filters were used (Whatman Anotop, 0.2 µm pore size, 10 mm diameter, Merck, Germany). Typical working solutions were prepared at a 5 mM total lipid concentration. Following preparation, liposomes were stored at 4 °C under N_2_ gas. All liposomes were characterized by size, particle concentration, and polydispersity index (PDI) by dynamic light scattering (Zetasizer ZS, Malvern, Panalytical, Malvern, UK).

To characterize the encapsulation efficiency of AuDDT, short-wave infrared (SWIR) fluorescence spectroscopy was performed using a SWIR-spectrometer from Wasatch (870–1700 nm; In-GaAs Hamamatsu 512 pixels; 6 nm resolution; f = 1.3; high pass filter at 830 nm) coupled with an optical fiber (*ø* = 300 μm, NA = 0.39) and using an 808 nm laser (120 mW cm^-2^) as excitation source. Samples at working concentration of 5 mM were measured in a quartz microcuvette (12.5 × 4.5 × 45 mm) with a 2 mm light path and spectra were recorded in the Wasatch software at high gain and 1000 ms acquisition time (average of 10 scans). Encapsulation efficiency was quantified by solving a standard curve based on the integrated fluorescence intensity (area under the curve 886–1703 nm) compared to the fluorescence intensity of loaded liposomes.

### Cryo-Electron Microscopy

Liposomes with a formulation of DOPE:DSPC:DSPE-PEG: Cholesterol (24:24:4:48 mol%) were prepared as described before, and concentrated by centrifugal ultrafiltration for 45 min at 3000 rcf using 15 mL Amicon cellulose filters (30 kDa MWCO, Sigma Aldrich, France). Liposomes at 40 mM final lipid concentrations were stored at 4 °C after purged with nitrogen gas (Messer CANgas, Sigma-Aldrich). Cryo-EM were prepared by applying 3 μL to glow-discharged 1.2/1.3-hole sized C-Flat holey copper grids with a carbon foil (300 mesh) and plunged frozen in liquid ethane using a Vitrobot Mark IV (Thermo Fisher Scientific, Waltham, MA, USA) (4 s blot time, blot force - 14). The sample was observed at the beamline CM01 of the European Synchrotron Radiation Facility (Grenoble, France [72]) with a Titan KriosG3 (Thermo Fisher Scientific) operating at 300 kV equipped with an energy filter (Bioquantum LS/967, AMETEK) (slit width of 20 eV). Images were recorded at a nominal magnification of 105000X (pixel size of 0.84 Å) on a K3 direct electron detector (AMETEK) in counting mode with EPU (Thermo Fisher Scientific). For image analysis, more than 500 images were selected for liposomes containing either 0, 0.2, 0.5, and 2 mol% AuDDT. For each subset, the liposomes were first classified by morphology, being unilamellar, multilamellar, or fused. Then the occurrence of gold nanoclusters was counted, both across the entire liposome population and within each morphological category. Smaller nanoclusters were counted individually, while aggregated clusters were counted as larger units based on their size and aggregation state.

Cryo-Electron Tomography was performed at the CM01 beamline of the European Synchrotron Radiation Facility [72]. The grids of 0.5 mol% AuDDT prepared for cryo-EM, were re-imaged by cryo-electron tomography. Tilt series were collected at a magnification of 33000X in super resolution counting mode corresponding to a pixel size of 1.38 Å. Movies were aligned using Motioncor2 and binned twice yielding a final pixel size of 2.76 Å. Alignment of tilt series and tomogram reconstruction were performed using AreTomo within the image processing pipeline for electron cryo-tomography in RELION-5.

### X Ray irradiation

X-ray irradiations were conducted using a Small Animal Radiation Research Platform (SARRP, Xstrahl, Walsall, UK). The beam was filtered through a combination of 0.8 mm of beryllium and 0.15 mm of copper and operated at 220 kVp and 13 mA. The focal source-to-sample distance was set at 35 cm, ensuring uniform exposure across the irradiation field. Samples were carefully positioned within a homogeneous dose distribution zone. Radiation was delivered at a mean dose rate of 3.06 Gy min^-1^ through a polyenergetic spectrum with an average photon energy of 78.54 keV.

### Detection of ROS production by X-rays

ROS generation by X-ray irradiations was assessed using the hydroxyl radical sensor aminophenyl fluorescein (APF, Merck). Reaction mixtures of 100 μL were prepared on black 96-well clear flat-bottom black plates (Greiner Bio-One, Frickenhausen, Germany), containing 10 μM APF and various liposomes at a final lipid concentration of 500 μM. Plates were irradiated at different doses of 2, 4, 8 and 16 Gy. The fluorescence of the reaction mixtures was measured before and after X-ray irradiation at *λ*_exc._ 485 ± 10 nm, *λ*_em._ 520 ± 20 nm (FLUOstar Omega, BMG Labtech, Ortenberg, Germany). All data was corrected to the background fluorescence measured at 0 Gy.

### Evaluation of cargo release induced by X-rays

To assess the capacity of radioresponsive-liposomes to release intraliposomal cargo, various AuDDT and AuDDT+BPD liposome compositions were loaded with a self-quenching solution of calcein (53 mM calcein, 180 mM NaCl, pH 7.4, 0.282 osmol/kg). After the sonication process, size-exclusion chromatography (SEC) using Sephadex G-50 fine columns hydrated with PBS were used to remove the non-encapsulated calcein [73]. Reaction mixtures of 100 μL were prepared, in which the various liposomes were diluted to a final concentration of 500 μM in PBS, fetal bovine serum (FBS), or Dulbecco’s Modified Eagle Medium (DMEM) supplemented with 10% FBS. The calcein emission was measured before and after X-ray irradiation and after lysis in 0.1% Triton X-100 (final concentration) at *λ*_exc._ 485 ± 10 nm, *λ*_em._ 520 ± 20 nm. All data was normalized to the maximum calcein emission following Triton X-100 addition.

### In vitro effects of radiotherapy responsive liposomes

Human pancreatic cancer cell lines PANC-1 (CRL-1469) and MIA PaCa-2 (CRL-1420) were obtained from the American Type Culture Collection (ATCC) and used between passages 2 and 30. Human pancreatic stellate cells (HPSCs) were purchased from ScienCell (San Diego, CA) and utilized between passages 2 and 15. Pancreatic cancer cells were cultured in DMEM supplemented with 5 mM GlutaMAX, 10% FBS, and 1% penicillin-streptomycin (10 000 Units/mL penicillin and 10 000 µg/mL streptomycin, Thermo Fisher Scientific). HPSCs were maintained in a 1:1 mixture of stellate cell-specific medium and supplemented DMEM. All cells were cultured under standard conditions (37°C, 5% CO₂, humidified atmosphere). Cell cultures were routinely evaluated for mycoplasma infection with MycoAlert® (Lonza, Basel, Switzerland) and were discarded when positive.

3D monoculture models, were established in 24-well plates (Greiner Bio-One, Kremsmünster, Austria) by embedding 7 500 pancreatic cancer cells per well in 220 µL of Matrigel (Corning, Corning, NY, USA). After 4 days of growth, microtumors were incubated overnight with liposomal formulations containing 0.5 mol% AuDDT at concentrations ranging from 100 µM to 2000 µM. The following day, microtumors were irradiated with single doses of 4 Gy, 8 Gy, or 16 Gy. Cultures were maintained until day 10, with media (containing 2% Matrigel) refreshed every two days.

3D heterotypic co-culture model was generated by co-seeding 5 000 pancreatic cancer cells with 5 000 HPSCs in 96-well U-bottom ultra-low attachment plates (Corning, NY, USA). Compact spheroids formed within 24 hours. On day 3 post-seeding, spheroids were incubated for 4 hours with one of the following liposomal formulations (final concentration 500 µM): empty liposomes, AuDDT (0.5 mol%), BPD (0.8 mol%), or combined AuDDT + BPD. Spheroids were subsequently irradiated with 4 Gy or 16 Gy of X-rays. Treatments were monitored up to day 14.

### Evaluation of Treatment Efficacy

At the experimental endpoints (day 10 for Matrigel microtumors, day 14 for heterotypic spheroids), treatment response was assessed using live/dead staining. Microtumors or spheroids were exposed to a solution containing 2 μM calcein AM (Invitrogen, Thermo Fisher Scientific) and 3µM propidium iodine (Sigma Aldrich) for 1h. Cellular viability was evaluated using confocal laser scanning microscopy (MICA microhub, Leica), using 20x objective (HC PL APO 20x/0.75) at λ_exc._ 488 nm, λ_em._ 495 – 540 nm and λ_exc._ 561 nm, λ_em._ 600 – 650 nm, respectively. Quantitative image analysis was conducted as previously described [74,75].

### In Vivo Biodistribution

All animal experiments were in accordance with the application for authorization of animal experiments submitted to the Ministry of Research and the Ethics Committee (APAFIS #33137-2021110411585349 v2). Female, 6-week-old NMRI nude mice (Janvier Labs) underwent surgery under isoflurane anesthesia to expose the spleen and pancreas. Subsequently, 1 × 106 PANC-1-pGL4 cells, suspended in 50 μL of Matrigel (Corning) were injected in the pancreas. Clinical follow-up consisted of behavior observation and monitoring of animal weight, performed 2–3 times a week. On day 24 after implantation, tumor size was assessed and confirmed by ultrasound imaging, animals were evenly distributed in treatment groups according to tumor size. AuDDT+BPD-PC liposomes containing 0.8 mol% 18:0 BPD-PC and 2 mol% AuDDT were prepared at a lipid concentration of 25 mM and a final BPD concentration of 200 μM, which was selected based on previous work [20]. Following randomization on day 35 post-implantation, once tumors reached approximately 100 mm^3^, mice bearing orthotopic PDAC received 200 μL (0.36 mg BPD/kg) of the AuDDT+BPD-PC liposomal solution via tail-vein injection under isoflurane anesthesia.

In vivo, whole-body fluorescence imaging (right and left sides, prone and supine positions) was performed prior to injection, and at 30 min, 90 min, 3 h, 5 h (6 mice), and 24 h (3 mice) post-injection under isoflurane anesthesia (Pearl Trilogy, LI-COR Biosciences GmbH, Bad Homburg, Germany) using *λ*exc. 685 nm, *λ*_em._ 700 – 720 nm. Animals were additionally imaged using 3D fluorescence tomography using a custom-built setup, followed by 3D CT (VivaCT, Scanco Medical, Wangen–Brüttisellen, Switzerland). Animals were humanely euthanized by cervical dislocation under gas anesthesia after 5 or 24 h post-injection. Organs were isolated and separately imaged using bioluminescence and fluorescence imaging (Li-Cor Pearl Trilogy) to quantify the distribution of BPD and BPD-PC. To image the biodistribution of the AuDDT delivered via AuDDT+BPD-PC liposomes, organs were imaged ex vivo using a custom-built shortwave infrared imaging setup (InGaAs Nirvana 640ST camera, Princeton), a 50 mm lens from Navitar (NA = 1.4), and a long pass filter at 1064 nm (Semrock), and excitation by 808 nm laser (120 mW cm^−2^) with a 200 ms acquisition time.

### Pharmacokinetics

Blood pharmacokinetics were assessed in healthy female 6-week-old NMRI nude mice (Janvier Labs). A single intravenous injection of 200 µL of the liposomal solution Au+BPD-PC (25mM) was administered via the tail vein (n = 3 mice per group). Blood samples (∼50 µL) were collected into lithium heparin-coated tubes at the following time points post-injection: 1 min, 5 min, 10 min, 20 min, 30 min, 3 h, 5 h, 24 h, and 54 h. At each time point, 30 µL of the collected blood was used for fluorescence-based quantification. Plasma (10 µL) was obtained by centrifugation and subsequently analyzed using fluorescence imaging. Two distinct fluorescence imaging modalities were employed to detect and quantify the two optical probes: NIR-I imaging (λ_exc_. 685 nm, λ_em._ 700 – 720 nm bandpass filter) for the detection of BPD, and SWIR imaging (λ_exc._ 808 nm, λ_em._ long-pass filter >900 nm) for the detection of AuDDT. Fluorescence background levels were established from plasma samples of non-injected control mice. All samples were imaged under identical settings, and the fluorescence signal was quantified using image analysis software (Li-Cor Pearl Trilogy). Results were expressed as relative light units per pixel (RLU/pixel), and data were reported as mean ± SEM for each time point (n = 3 mice/group). Fluorescence intensity curves over time were plotted to assess blood circulation profiles. The elimination half-life (t₁/₂) of the fluorescent liposomes was calculated using non-linear regression analysis in GraphPad Prism 7 (GraphPad software, San Diego, CA, USA), based on a mono- or bi-exponential decay model depending on the fit quality.

### Inductively-coupled plasma mass spectrometry (ICP-MS)

Liposomal formulations composed of DOPE:DSPE-PEG: Cholesterol (48:4:48 mol%) containing increasing concentrations of AuDDT were prepared as previously described. Afterward, a known volume of the prepared formulation was dried using vacuum centrifugation (Genevac mi-Vac centrifugal concentrator, 40 °C, 24 h minimum) to get liposome pellets. Thereafter, aqua regia (1:3 molar ratio of HNO3 (67% wt, Merck, Germany) and HCl (35% wt, Merck, Germany), respectively) was added to the pellets and left for overnight incubation. On the next day, the digestion was completed through heating at 70 °C for spaced periods of 10 minutes for a total of 1h. Before the ICP-MS measurements, the digested samples were diluted with deionized water to a final volume of 10 mL and transferred to the ICP-MS tubes. A calibration curve of gold concentrations up to 1000 ppb (μg L^−1^) was obtained using the analytical gold standard TraceCERT, 1 g L^−1^ Au in hydrochloric acid (Merck, Germany). The NexION 1000 ICP-MS instrument (Perkin Elmer, Shelton, CT, USA) was used for the measurements.

In parallel, *ex vivo* quantification of gold biodistribution was performed on tissue samples collected from tumor, liver, and spleen at various post-injection time points. Immediately after collection, tissues were weighed and processed using the same digestion and dilution protocol as described for the liposomal formulations. This allowed direct comparison of gold accumulation across organs and over time following systemic administration of AuDDT-containing liposomes.

### Laser-induced breakdown spectroscopy (LIBS)

LIBS elemental imaging was performed on ex vivo tumor cryosections (10 μm) analyzed at room temperature, with ELM-XS MED (Ablatom S.A.S, Villeurbanne, FR) using a setup previously described [76,77]. Briefly, a Nd: YAG laser (1064 nm, pulse duration = 8 ns, repetition rate = 100 Hz) was focused on the target through a microscope objective. Samples were placed on a 3D motorized stage and were scanned pixel by pixel, with a lateral resolution of 30 µm and an energy of 6 mJ per pulse. Each pixel was a result of a single shot and no average or accumulation was performed. A flow of 0.8 L min^−1^ of argon gas was directed on the plasma region during the whole experiment. The first spectrometer collected the plasma light through an entrance slit of 90 µm and a grating of 2400 lines mm^−1^ (240 nm blaze), to ensure a high resolution in the UV for detecting intense line of phosphorus at 253.4 nm and gold at 242.7 nm. The delay and gate detection were set to 3 µs, with a gain of 1000. The acquisition and data analysis were performed using a custom-developed LabVIEW software (National Instruments, Austin TX, USA).

### Statistical Analyses

All data was statistically analyzed in Graphpad Prism 7 (Graphpad, USA.). Data sets were tested for Gaussian distributions using D’Agostino and Pearson omnibus normality tests. Normally distributed data sets were analyzed using a One-way ANOVA and Tukey’s post-hoc test for multiple comparisons, whereas non-Gaussian data sets were analyzed using a Kruskal-Wallis for multiple comparison. For ROS production and liposome-cargo release assays, data were fitted using dose-response fits (agonist vs response (three parameters)), and statistically compared using an extra-sum-of-squares F-test. For tumor growth comparisons, non-linear regression of logistic growth was applied, regarding dose-response assays curve fitting was performed using a nonlinear regression model (inhibitor vs. normalized response, variable slope). Cleaned data were processed by removing outliers using the ROUT method (Q = 10%). Statistical significance is indicated with single asterisks (p≤0.05), double asterisks (p≤0.01) or triple asterisks (p≤0.005) or quadruple asterisks (p≤0.001).

## Acknowledgments

This research was supported by the French Institute for Health and Medical research (INSERM), and the European Union M.B: (ERC STG, RADIOCONTROL, 101078392). Views and opinions expressed were, however, those of the authors only and did not necessarily reflect those of the European Union or the European Research Council Executive Agency. Neither the European Union nor the granting authority could be held responsible for them. X.L.G. gratefully acknowledges funding by the Agence National de la Recherche NanoGOLD (ANR-22-CE29-0022). The authors additionally acknowledge the European Synchrotron Radiation Facility for the provision of beam time on CM01 (inhouse research). Grenoble IRMaGe facility was partly funded by the French program “Investissement d’Avenir” run by the ‘Agence Nationale pour la Recherche’; grant ‘Infrastructure d’avenir en Biologie Santé’ -ANR-11-INBS-0006. The SAXO equipment (SARRP) was partly funded by the ITMO Cancer of Aviesan within the framework of the 2021-2030. Cancer Control Strategy, on funds administrated by INSERM”; The rotary club “Jetons le cancer 2022”; France Life Imaging, grant “ANR-11-INBS-0006”; The Grenoble Alpes University (UGA); the GEFLUC (2022) and the Labex PRIMES ANR-11-LABX-0063, The CALYPSO project, INSERM Cancer PCSI 20CP082-00.

