## Supplemental information for "Radiotherapy Enhancement by Gold Nanocluster-functionalized Nanoliposomes Using Polychromatic Orthovoltage X-ray Irradiation"

*Corresponding author

Mans Broekgaarden

Institute for Advanced Biosciences

Université de Grenoble-Alpes, Inserm U1209, CNRS UMR 5309

Allée des Alpes

38700 La Tronche

France

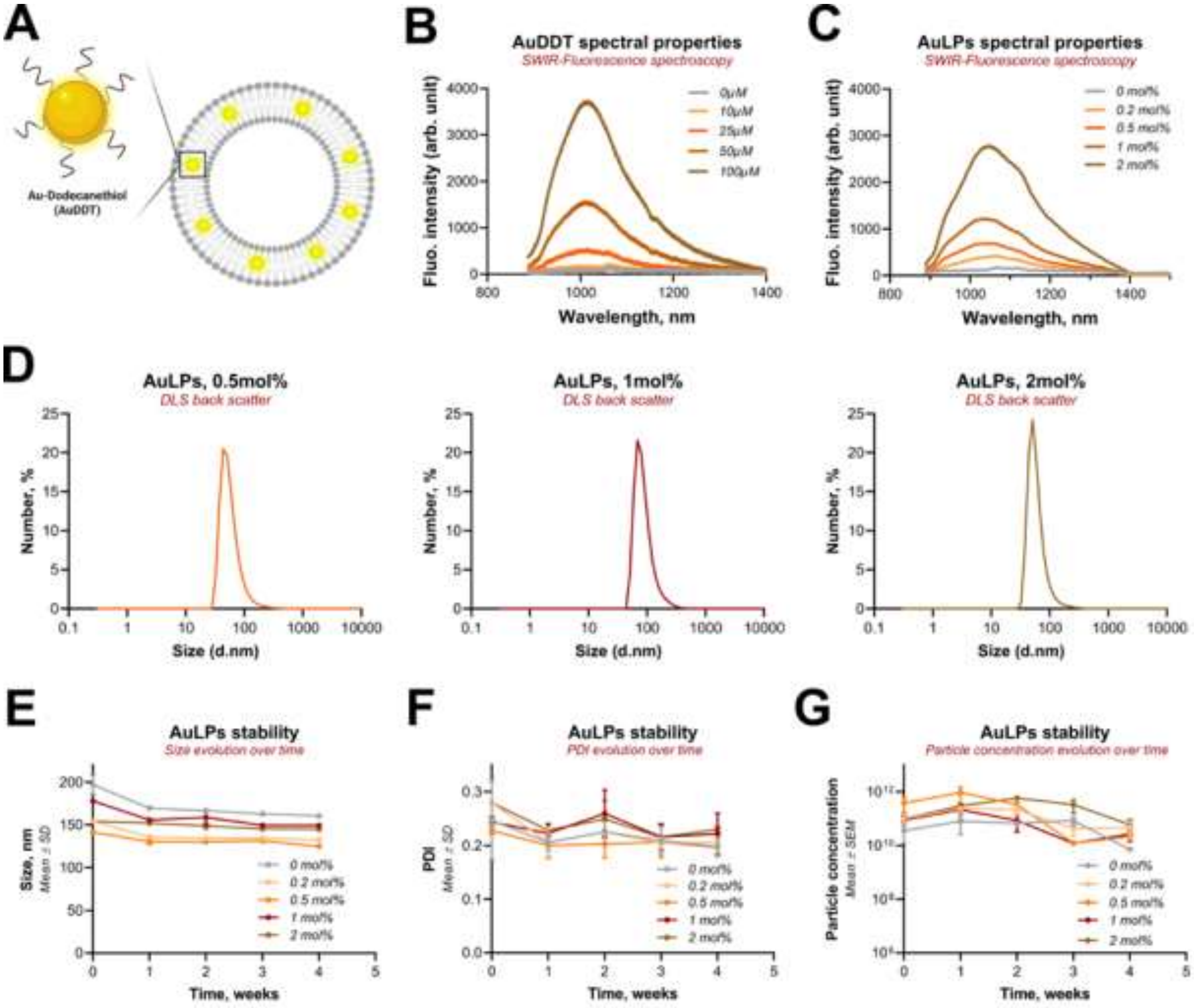


**Figure S1. AuLPs characterization and colloidal stability.** (A) Schematic representation of AuDDT cluster embedded in the liposomal lipid bilayer. (B) SWIR-fluorescence spectra of AuDDT nanoclusters in chloroform at increasing concentrations (0 - 100µM). (C) SWIR-fluorescence spectra of AuLPs in PBS at increasing and corresponding concentrations of AuDDT (0 – 2mol%). (D) Size distribution of freshly prepared AuLPs containing increasing concentrations of AuDDT (0.5–2 mol%) measured by DLS. (E) Evolution of liposome size, PDI (F) and particle concentration (G) over one month of storage at 4 °C. For G-I, the data represents n = 5-15 from three independent batches.

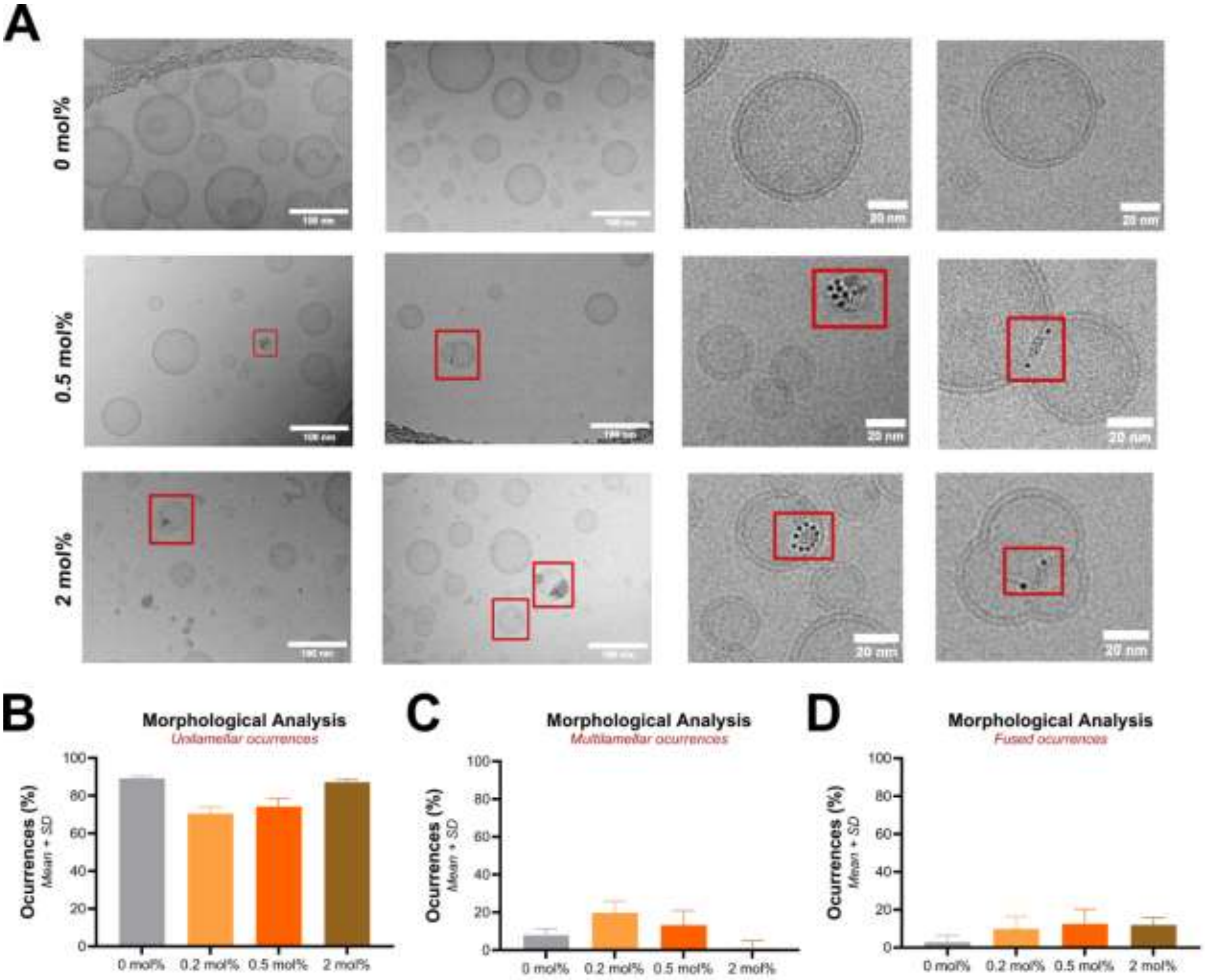


**Figure S2. Au LPs morphological characterization through cryo-electron microscopy.** (A) Representative cryo-EM images of liposomes loaded with 0, 0.5 mol% or 2 mol% of AuDDT zoomed-in areas, distinct electron-dense spherical feature embedded in the lipid bilayer are identified as AuDDT nanoclusters, pointed within red frame (scale bar = 100nm and 20nm). Morphological classification of vesicle lamellarity across all formulations (0, 0.5, 1, and 2 mol% AuDDT) indicating the percentage of liposomes founded as unilamellar (B), multilamellar (C) or fused (D) vesicles.

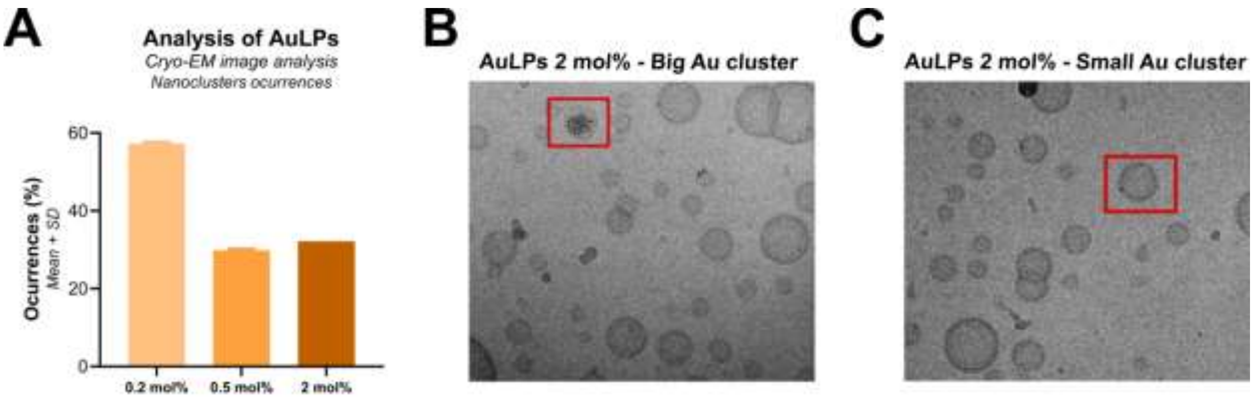


**Figure S3. Au LPs cluster analysis and quantification.** (A) Cluster occurrences in all liposomal formulations 0.2, 0.5, and 2 mol% loaded with AuDDT observed by cryo-EM. Representative liposomal structures containing big AuDDT clusters (B) and small and unitarian AuDDT clusters (C) embedded on liposomal formulations, highlighted by red frames.

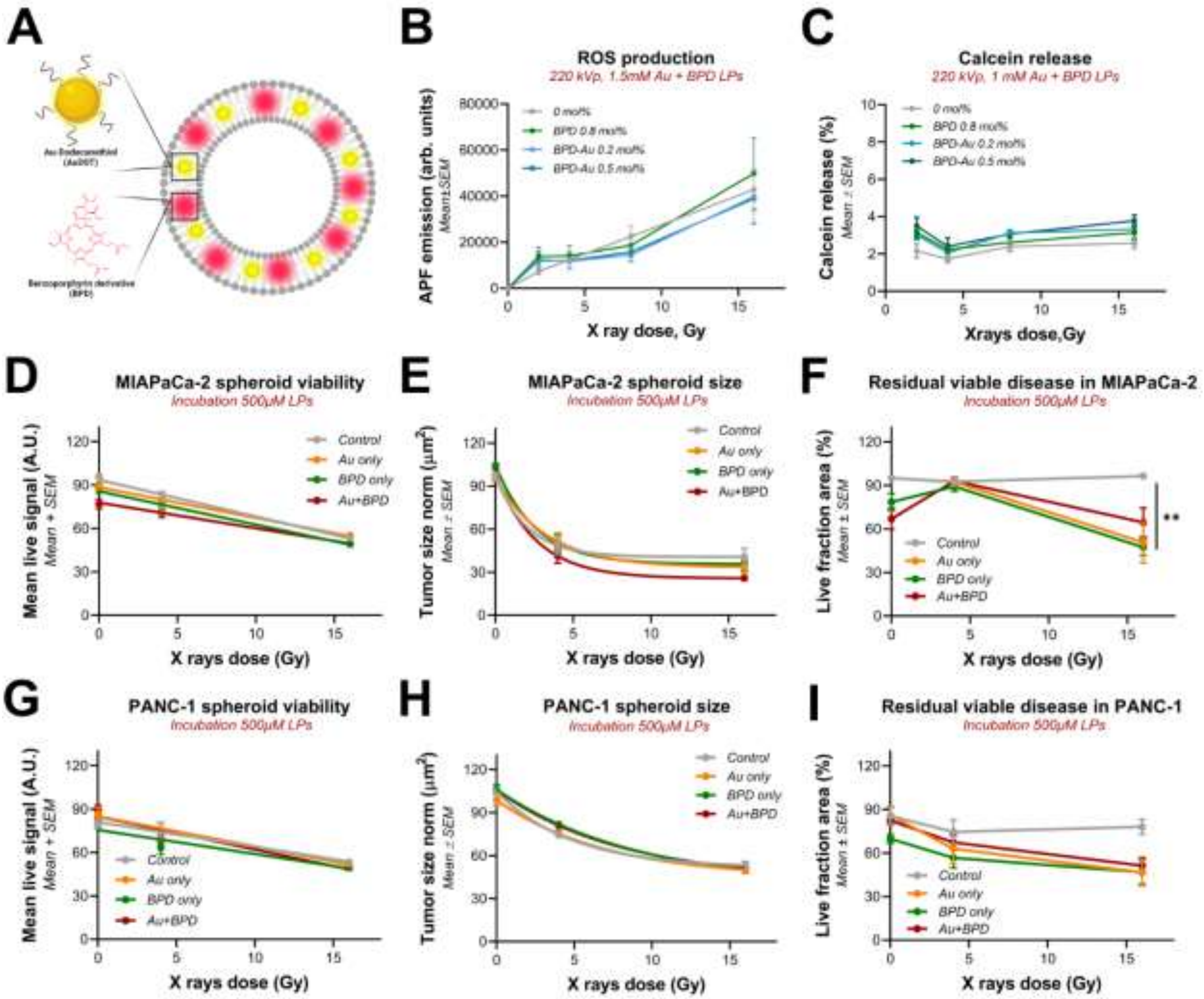


**Figure S4. Performance evaluation of Au+BPDLPs for ROS production, drug release and *in vitro* efficacy.** (A) Schematic representation Au+BPD LPs with AuDDT and BPD integrated in the lipid bilayer. (B) ROS production measured by APF oxidation under increasing doses of X-ray irradiation in different formulations of Au+BPD LPs loaded with 0.2 or 0.5 mol% of AuDDT with or without 0.8mol% BPD. (C) Immediate calcein release from Au+BPD LPs loaded with 0.2 or 0.5 mol% of AuDDT with or without 0.8mol% BPD. For B-C, the data represent n = 18-25 from three independent technical repetitions. *In vitro* efficacy of non-loaded (gray), AuLPs (orange), BPD LPs (green) and Au+BPD LPs (red) on MIAPaCa-2 (D-F) and PANC-1 (G-I) spheroids. (D and G) Viability quantification, as mean live signal, across all treatments and irradiation doses. (E and H) Normalized spheroid size quantification across all treatments and irradiation doses. (F and I) Residual viable disease, as live fraction area, across all treatments and irradiation doses. Data are presented as means from three independent replicates per experiment, with sample sizes of N = 6 - 18. Data were not normally distributed and analyzed using one-way ANOVA with Kruskal-Wallis multiple comparison test.

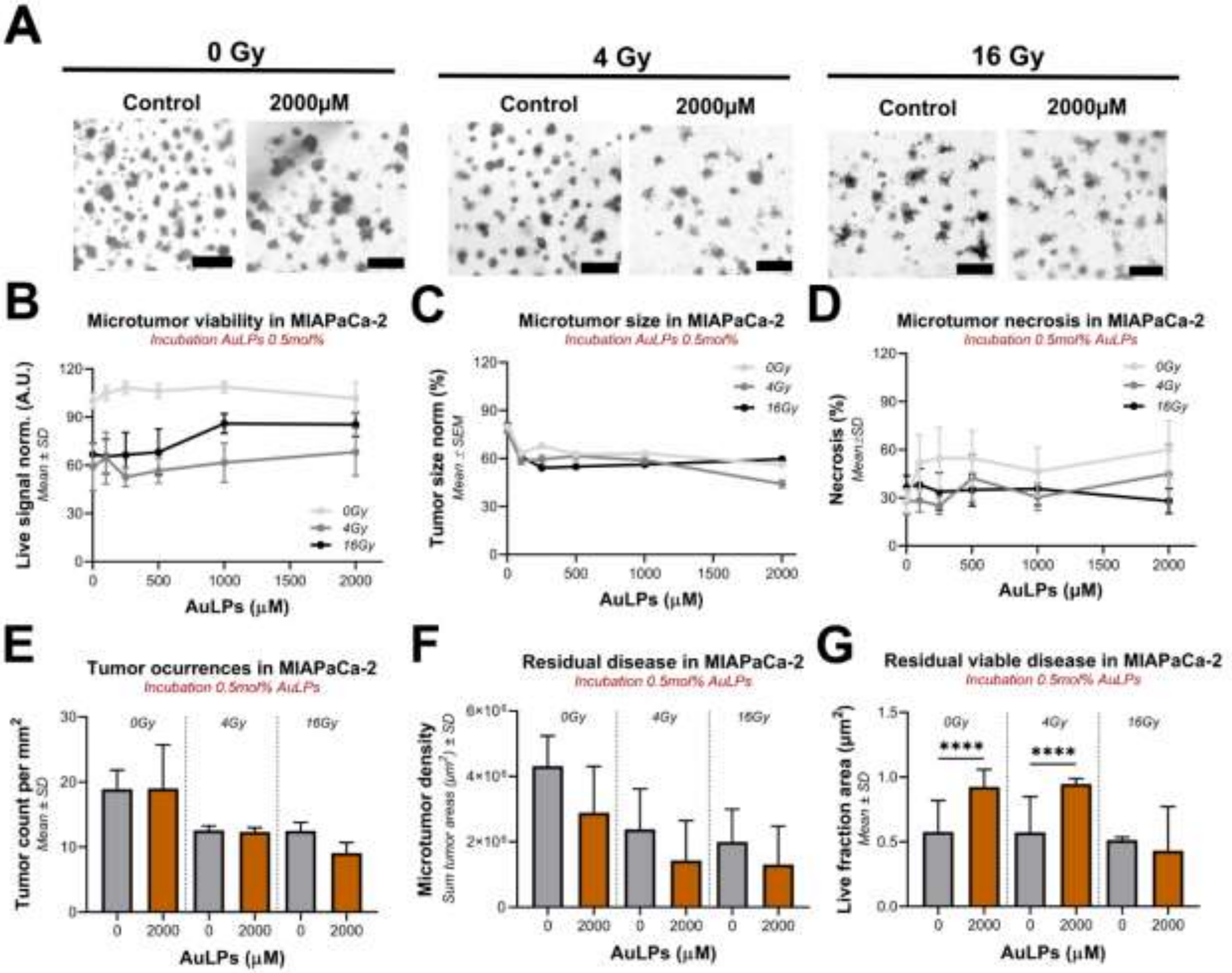


**Figure S5. Effect of AuLPs (0.5 mol%) and X-ray irradiation on 3D MIAPaCa-2 microtumors**. (A) Brightfield images of microtumors without irradiation (0 Gy) and at 4 Gy and 16 Gy doses, scale bar 400µm. (B) Quantification of microtumor viability in relation to liposome concentration for MIAPaCa-2 microtumors at different Xrays doses, 0, 4 and 16 Gy. (C) Normalized microtumor size quantification in relation to liposome concentration for MIAPaCa-2 microtumors at different Xrays doses, 0, 4 and 16 Gy. (D) Normalized microtumor necrosis quantification in relation to liposome concentration for MIAPaCa-2 microtumors at different Xrays doses, 0, 4 and 16 Gy. (E) Tumor occurrences per mm$^2$ for non-loaded AuLPs (in grey) and 0.5mol% AuLPs at 2mM concentration (in dark orange) across the different irradiation doses. (F) Residual disease as the sum of microtumor areas per condition, for non-loaded AuLPs (in grey) and 0.5mol% AuLPs at 2mM concentration (in dark orange) across the different irradiation doses. (G) Residual viable disease as the live fraction area remaining per condition, for non-loaded AuLPs (in grey) and 0.5mol% AuLPs at 2mM concentration (in dark orange) across the different irradiation doses. Data are presented as means from three independent replicates per experiment, with sample sizes of n = 53-1750.

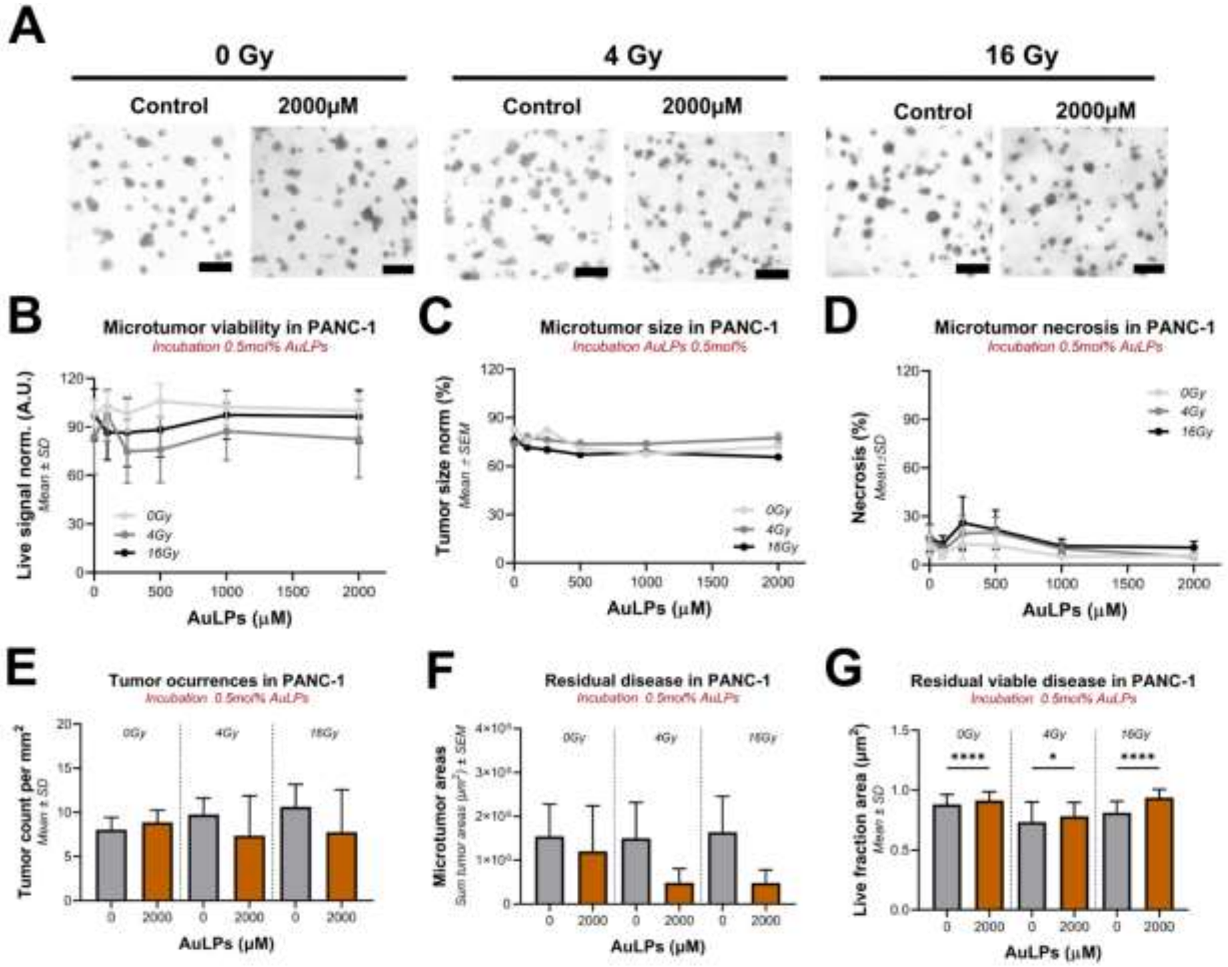


**Figure S6. Effect of AuLPs (0.5 mol%) and X-ray irradiation on 3D PANC-1 microtumors.** (A) Brightfield images of microtumors without irradiation (0 Gy) and at 4 Gy and 16 Gy doses, scale bar 400µm. (B) Quantification of microtumor viability in relation to liposome concentration for PANC-1 microtumors at different Xrays doses, 0, 4 and 16 Gy. (C) Normalized microtumor size quantification in relation to liposome concentration for PANC-1 microtumors at different Xrays doses, 0, 4 and 16 Gy. (D) Normalized microtumor necrosis quantification in relation to liposome concentration for PANC-1 microtumors at different Xrays doses, 0, 4 and 16 Gy. (E) Tumor occurrences per mm$^2$ for non-loaded AuLPs (in grey) and 0.5mol% AuLPs at 2mM concentration (in dark orange) across the different irradiation doses. (F) Residual disease as the sum of microtumor areas per condition, for non-loaded AuLPs (in grey) and 0.5mol% AuLPs at 2mM concentration (in dark orange) across the different irradiation doses. (G) Residual viable disease as the live fraction area remaining per condition, for non-loaded AuLPs (in grey) and 0.5mol% AuLPs at 2mM concentration (in dark orange) across the different irradiation doses. Data are presented as means from three independent replicates per experiment, with sample sizes of n = 64-1107.

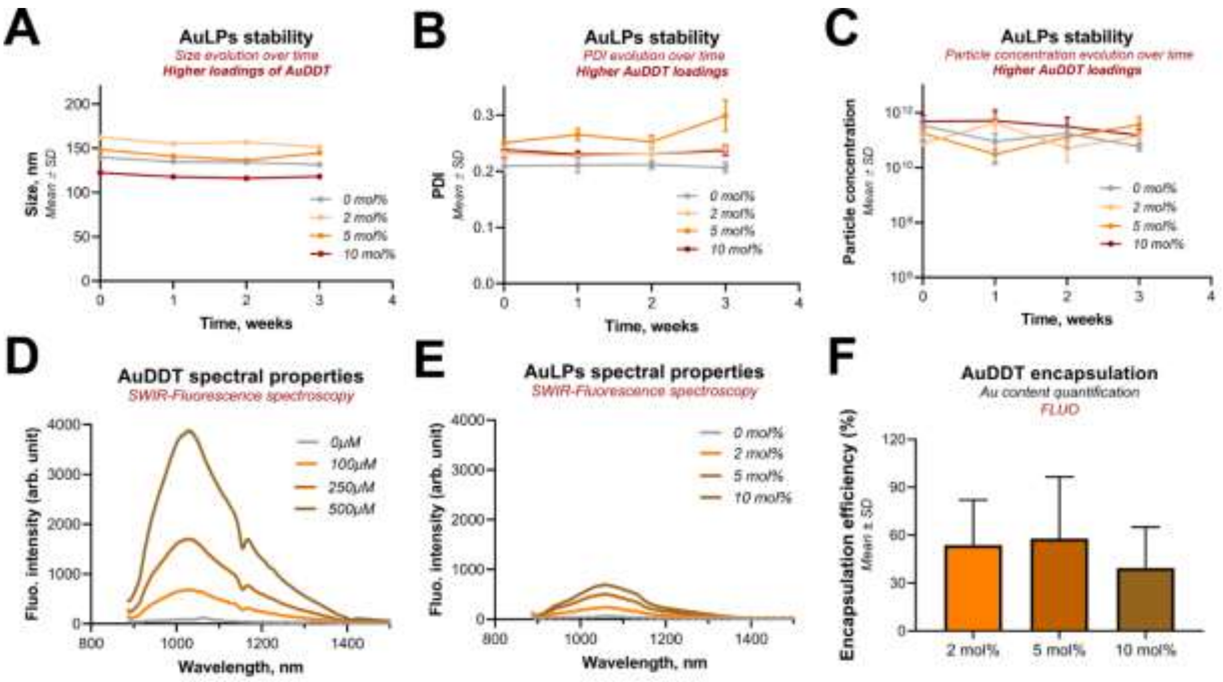


**Figure S7. Stability over time of higher loadings of AuDDT LPs**. (A) Evolution of liposome size, (B) PDI (C) and particle concentration over one month of storage at 4 °C. (D) SWIR-fluorescence spectra of AuDDT nanoclusters in chloroform at increasing concentrations (0 - 500µM). (E) SWIR-fluorescence spectra of AuLPs in PBS at increasing and corresponding concentrations of AuDDT (0 – 10mol%). (F) AuDDT semiquantitative encapsulation calculated as nanocluster fluorescence on AuLPs. For A to C, data represents n = 8, from D to F, data represents n=3.

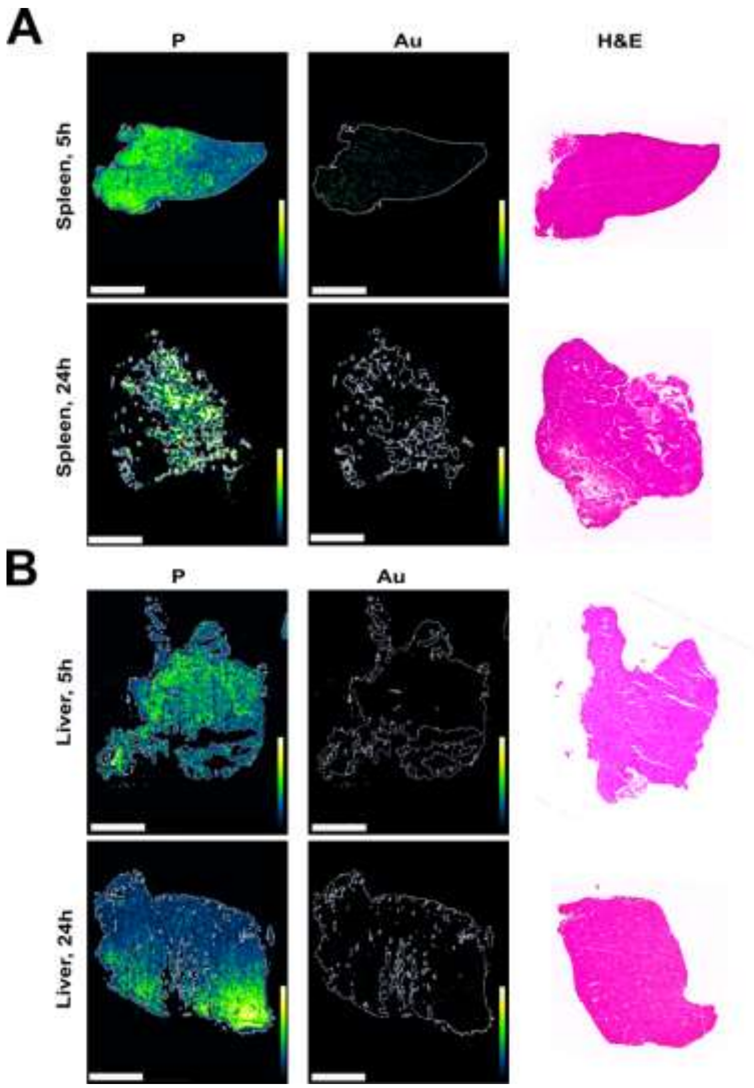


**Figure S8. Elemental distribution and histological evaluation of AuLPs in tumor-bearing mice.** (A) Elemental mapping of orthotopic PANC-1 tumor sections by LIBS on Au+BPD LPs treated mice, depicted are spleen, (B) and liver 5- and 24-hours post-injection, quantification of phosphorus (P), and gold (Au) performed. Representative H&E staining of the tissue slide selected is also shown. Scale bar, 2 mm. Scale bar, 2 mm.

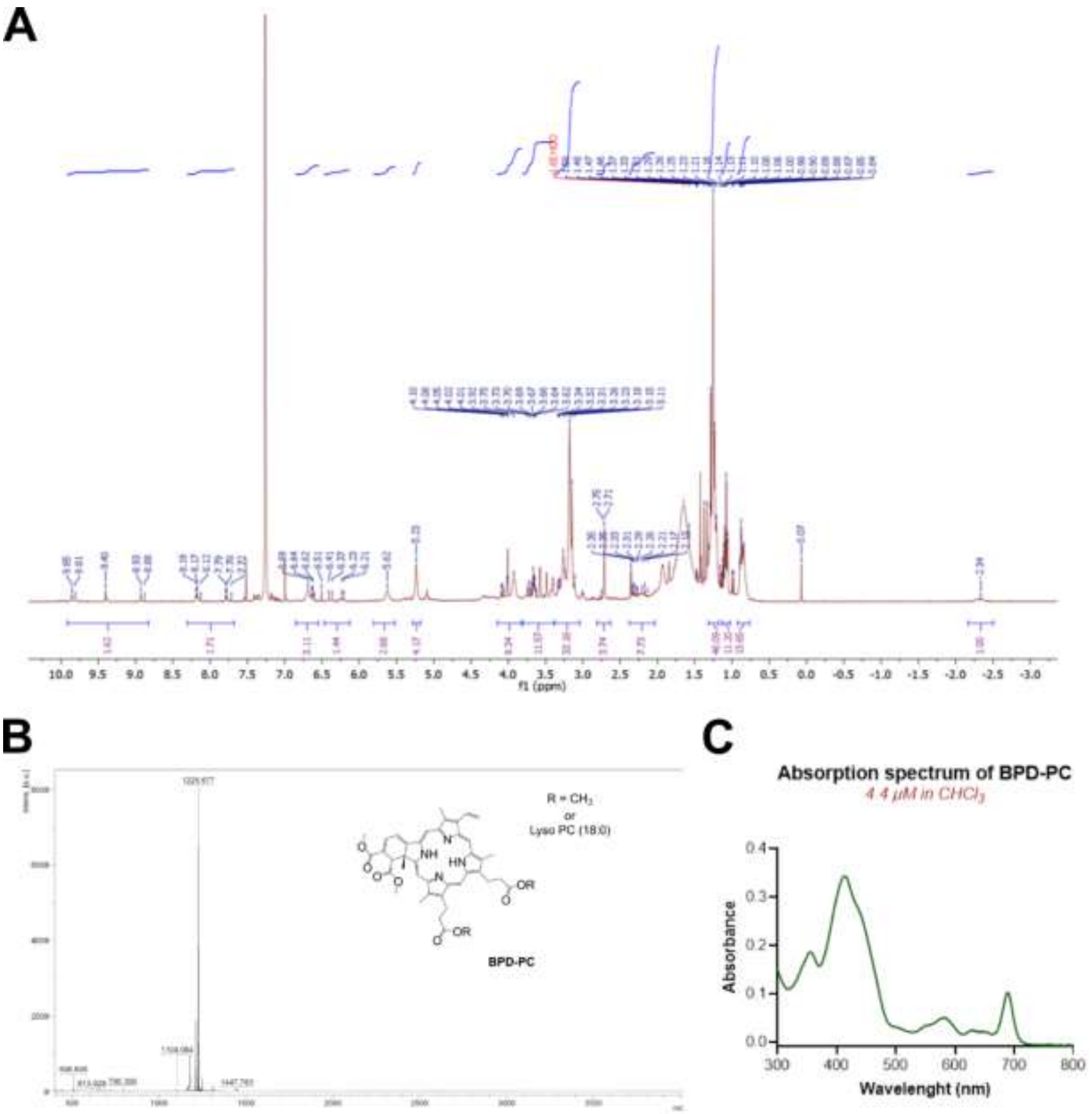


**Figure S9. Characterization of BPD-PC.** (A) $^{1}$H NMR spectra of purified BPD-PC $^{1}$H NMR (400 MHz, CDCl3-d, δ) 9.83 (d, *J* = 16.2 Hz, 1H), 8.32 – 7.98 (m, 1H), 6.88 – 6.57 (m, 2H), 6.45 – 5.99 (m, 1H), 5.62 (s, 2H), 5.16 (d, *J* = 57.5 Hz, 4H), 4.29 (dd, *J* = 13.6, 6.5 Hz, 2H), 4.15 – 3.82 (m, 5H), 3.80 – 3.47 (m, 2H), 3.16 (d, *J* = 13.3 Hz, 15H), 2.93 – 2.54 (m, 2H), 2.45 – 2.06 (m, 2H), 2.04 – 1.70 (m, 5H), 1.45 – 1.15 (m, 32H), 1.15 – 1.03 (m, 6H), 0.95 – 0.76 (m, 10H), -2.34 (s, $^{1}$H). (B) MALDI-TOF mass spectra of purified BPD-PC: m/z $[M+H]^+$ calculated for $C_{67}H_{94}N_5O_{14}P$, 1224.48; found = 1225.68. (C) UV – Vis Absorption spectra of purified BPD-PC in chloroform: $\lambda_{max}$ (ε) = 689 nm (23139), 581 nm (11526), 413 nm (77493), 356 nm (42187).
